# Melody: Decoding the Sequence Determinants of Locus Specific DNA Methylation Across Human Tissues

**DOI:** 10.1101/2025.11.23.689975

**Authors:** Junru Jin, Ding Wang, Jianbo Qiao, Wenjia Gao, Yuhang Liu, Siqi Chen, Quan Zou, Shu Wu, Ran Su, Leyi Wei

**Affiliations:** School of Software, Shandong University, Jinan, Shandong Province, China; New Laboratory of Pattern Recognition (NLPR), State Key Laboratory of Multimodal Artificial Intelligence Systems (MAIS), Institute of Automation, Chinese Academy of Sciences, Beijing, China; School of Artificial Intelligence, University of Chinese Academy of Sciences, Beijing, China; Faculty of Applied Sciences, Macao Polytechnic University, Macao SAR, China; Institute of Fundamental and Frontier Sciences, University of Electronic Science and Technology of China, Chengdu, Sichuan Province, China; College of Intelligence and Computing, Tianjin University, Tianjin, China; Engineering Research Centre of Applied Technology on Machine Translation and Artificial Intelligence, Macao Polytechnic University, Macao SAR, China

**Author notes:** Correspondence: Shu Wu, Ran Su, Leyi Wei. These authors contributed equally (Junru Jin, Ding Wang, Jianbo Qiao). These authors jointly supervised this work (Shu Wu, Ran Su, Leyi Wei).

## Abstract

DNA methylation is a fundamental epigenetic modification that plays crucial roles in transcriptional regulation, cellular differentiation, and genome stability. However, how locus-specific DNA methylation is determined by intrinsic DNA sequence remains poorly understood. Here, we introduce Melody, a deep learning framework that predicts DNA methylation from 10-kb genomic sequences, enabling the integration of both local and long-range sequence signals. Across 39 human tissues, Melody accurately predicts methylation profiles and consistently outperforms existing state-of-the-art methods in whole-chromosome, hypomethylated-region, and cell-type-specific benchmarks. Melody also generalizes to methylation quantitative trait locus (meQTL) effect prediction and identifies regulatory sequence motifs associated with methylation variability. To extend prediction beyond profiled tissues, we further develop Melody-G, which incorporates single-cell RNA-seq foundation model embeddings to infer methylation states in previously unseen cell types directly from transcriptomic data. Together, Melody provides a unified framework for linking genomic sequence and cellular state to DNA methylation and offers new insights into the regulatory logic governing the human methylome.

## Introduction

DNA methylation is a fundamental epigenetic modification that regulates gene expression, maintains genome stability, and contributes to tissue-specific cellular identity^1,2^. It plays critical roles in diverse biological processes such as embryonic development, cell differentiation, and aging. Aberrant DNA methylation patterns have been implicated in a wide range of diseases, including cancers, neurological disorders, and metabolic syndromes^3–5^. DNA methylation profiles have also been shown to reflect biological age and exposure history, making them a powerful resource for both basic research and clinical applications^6^. In addition, methylation Quantitative Trait Loci (meQTLs)^7^, genetic variants that influence DNA methylation levels, serve as important molecular markers for understanding disease mechanisms and have emerged as a valuable framework to study the relationship between genetic variation to DNA methylation and gene expression^8^.

With the growing availability of large-scale sequencing data, deep learning has emerged as a transformative approach for decoding the non-linear regulatory relationships encoded in the genome^9,10^. Models such as AlphaGenome^11^, Enformer^12^, have demonstrated remarkable capacity in capturing distal regulatory dependencies from raw DNA sequence. However, despite these advances, current frameworks, including AlphaGenome, do not explicitly model DNA methylation or its cell-type specificity. Several earlier studies have proposed specialized models for methylation prediction, such as DeepCpG^13^, CPGenie^14^, iDNA-ABT^15^, iDNA-ABF^16^ and INTERACT^17^, which use local genomic sequence context to predict methylation states. DeepCpG and CPGenie represent early convolutional architectures that rely on shallow two-layer CNNs to extract local sequence features associated with methylation. In contrast, iDNA-ABT and iDNA-ABF introduce transformer-based and pretrained architectures to capture sequence dependencies, while INTERACT integrates convolutional and transformer layers within a unified framework. While these pioneering efforts established the feasibility of deep learning for methylation analysis, their limitations have become increasingly apparent in light of recent methodological and data advances.

First, existing methylation training datasets and prediction frameworks have several important limitations. Many previous models were trained on limited or early-generation datasets and relied on short sequence windows (e.g., 41 bp), which fail to capture dependencies that are now known to influence methylation. Moreover, previous models were designed for a single or few cell lines, overlooking the extensive cell-type-specific heterogeneity of methylation across tissues. Second, meQTLs prediction, a key question in functional genomics, remains poorly benchmarked in the DNA methylation field. No standardized benchmark dataset exists to fairly evaluate model performance on meQTL effect prediction, leaving the field without a clear assessment of core DNA methylation motif capture. Third, most existing models exhibit poor generalizability: they can predict methylation accurately only on training datasets but fail to extend to new tissues or developmental stages. This lack of generalization severely limits their clinical and translational applicability, even though DNA methylation profiling is increasingly used in cancer diagnostics, disease screening, and biological age estimation.

To address these challenges, we developed Melody (Methylation Learning On DNA), a deep learning framework for decoding DNA methylation from sequence and transcriptomic information. Unlike previous models, Melody is built to bridge the gap between sequence and tissue-specific epigenome regulation. It provides a scalable solution for inferring DNA methylation profiles across diverse biological contexts, enabling the discovery of regulatory motifs and the interpretation of non-coding variants. Furthermore, we construct comprehensive meQTLs benchmark datasets to systematically evaluate variant effect prediction, providing fair and unified comparison among models. In addition, we perform a detailed analysis of cell-type-specific motifs, characterizing them in terms of both specificity and regulatory strength to uncover sequence determinants of methylation variation. We also provide a powerful tool Melody-G that improves performance on cell-type-specific regions using transcriptomic profiles. To achieve these functionalities, we implemented Melody through three specialized architectural variants: a multi-track Melody-MT predictor for shared cross-tissue feature learning and downstream generalization, a single-track Melody-ST predictor for cell-type-specific prediction, a generalized model Melody-G that integrates DNA sequence with single-cell RNA-seq based cell-state embeddings to infer methylation in unseen biological contexts. For accurate methylation modeling, we extended the input sequence window to 10 kb, which allows the model to encompass distal motifs and flanking CpG islands known to modulate local methylation states. To ensure robust performance, we also implement multi-task learning that combines CpG-level prediction with auxiliary regional tasks. To support broad accessibility, we also developed a user-friendly Melody web server (https://inner.wei-group.net/Melody/) for interactive prediction and visualization of methylation landscapes across tissues.

## Results

### Melody variants enable DNA methylation prediction across diverse contexts

Locus-specific CpG methylation varies sharply across the genome and across human cell types, but the extent to which these patterns are encoded in DNA sequence remains incompletely understood. We therefore curated methylation maps for 39 normal human cell types^18^ and trained Melody to predict continuous CpG methylation levels from 10-kb genomic sequence windows (Fig. 1A). Melody uses a fully convolutional encoder–decoder backbone to integrate sequence information across multiple genomic scales while retaining position-level methylation predictions. The primary output predicts methylation levels at CpG sites, and two auxiliary 100-bp tasks—regional methylation and CpG density—regularize the model to capture broader methylation architecture. This design contrasts with previous methylation predictors that use substantially shorter sequence windows and are therefore less able to capture distal sequence context.

**Figure 1.**
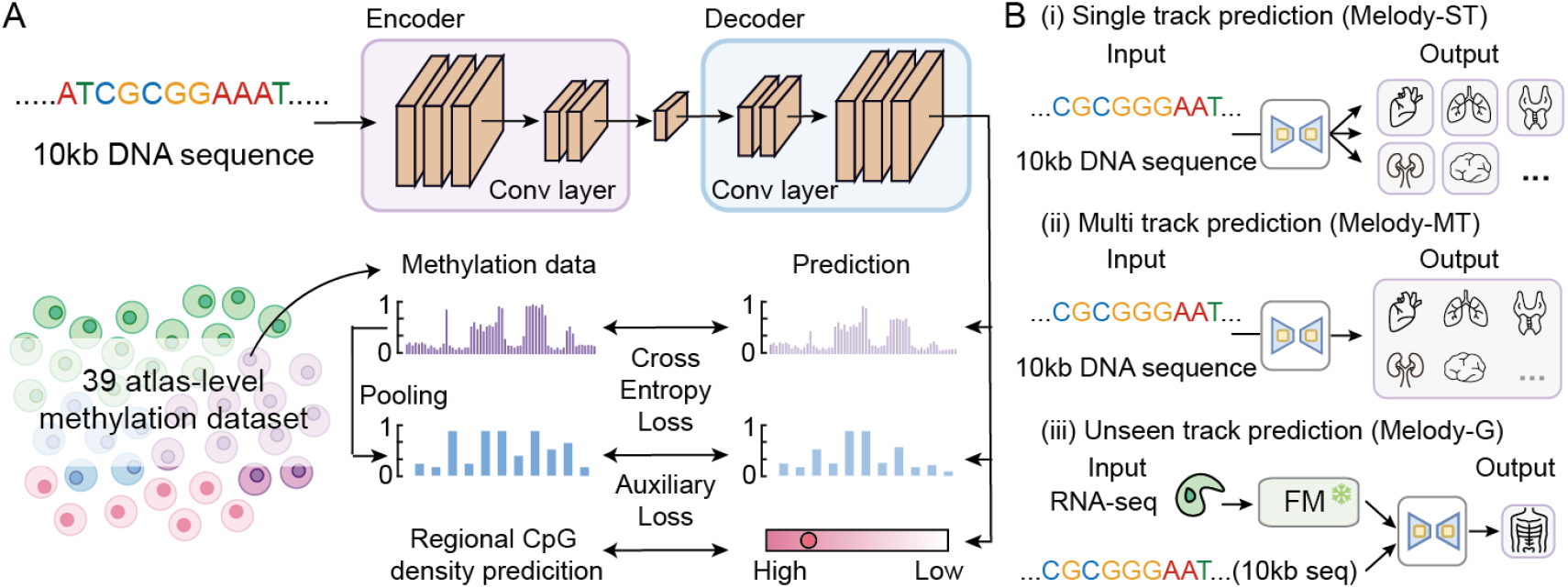
Overall framework and workflow of the Melody framework. (A) Melody takes 10 kb DNA sequence input and uses Encoder and Decoder composed of convolutional layers to capture DNA features. A multi-task prediction head subsequently performs joint learning for single-base resolution methylation levels, average methylation levels per 100 bp, and CpG counts per 100 bp. (B) Melody encompasses three variants, namely Melody-ST, Melody-MT, and Melody-G, which are optimized for distinct task scenarios.

We implemented three Melody configurations for complementary tasks, namely Melody- ST, Melody-MT, and Melody-G (Fig. 1B). Melody-ST and Melody-MT are tailored for single-track and multi-track methylation profiling respectively. While multi-track model can leverage shared information across cell types, prior studies^19^ have suggested that such models may perform less well in cell-type-specific regulatory regions, where modeling shared structure can come at the cost of cell-type resolution. As a result, training a single-track model for an individual cell type provides a complementary strategy for capturing more cell-type-specific methylation patterns. Melody-G incorporates cell-type-specific scRNA-seq data to enable methylation level prediction in unseen cell types guided by cellular identity (Supplementary Figure S1), which is a variant that integrates cell-type-specific scRNA-seq embeddings derived from pre-trained single-cell foundation models (FM). Melody-G employs a feature-wise linear modulation strategy to fuse these cellular features with DNA sequence representations, effectively conditioning the network to identify methylation states across diverse cellular contexts. We designed a cross-cell-type generalization experiment wherein the model was trained on data from a set of observed cell types and evaluated on data from a completely disjoint set of unobserved cell types. Melody-G undergoes a two-phase training regimen. In Phase 1 (G1), the model is trained on whole chromosomes from seen cell types. Subsequently, Phase 2 (G2) focuses on fine-tuning using cell-specific regions from seen cell types to facilitate generalization to specific regions in unseen cell types.

### Melody outperforms existing methods in methylation prediction

We benchmarked Melody against leading DNA methylation prediction methods, including INTERACT, CpGenie and iDNA-ABT, under three complementary evaluation regimes: chromosome-split validation, hypomethylated sample-split validation and cell-type-specific regions. Here, we focused on two Melody variants, Melody-ST and Melody-MT, designed to capture and interpret regulatory determinants of DNA methylation. Across 39 datasets, Melody-MT achieved mean Spearman correlations of 0.723 on the sampling test set and 0.645 on the test chromosome, outperforming the strongest competing baselines (0.590 and 0.584 on the corresponding sets)(Figure 2A,B, Supplementary Figure S2, S3). Melody-ST likewise improved upon all baselines across metrics, and together the two variants yielded an average ∼17% relative gain in Spearman rank correlation over competing methods.

**Figure 2.**
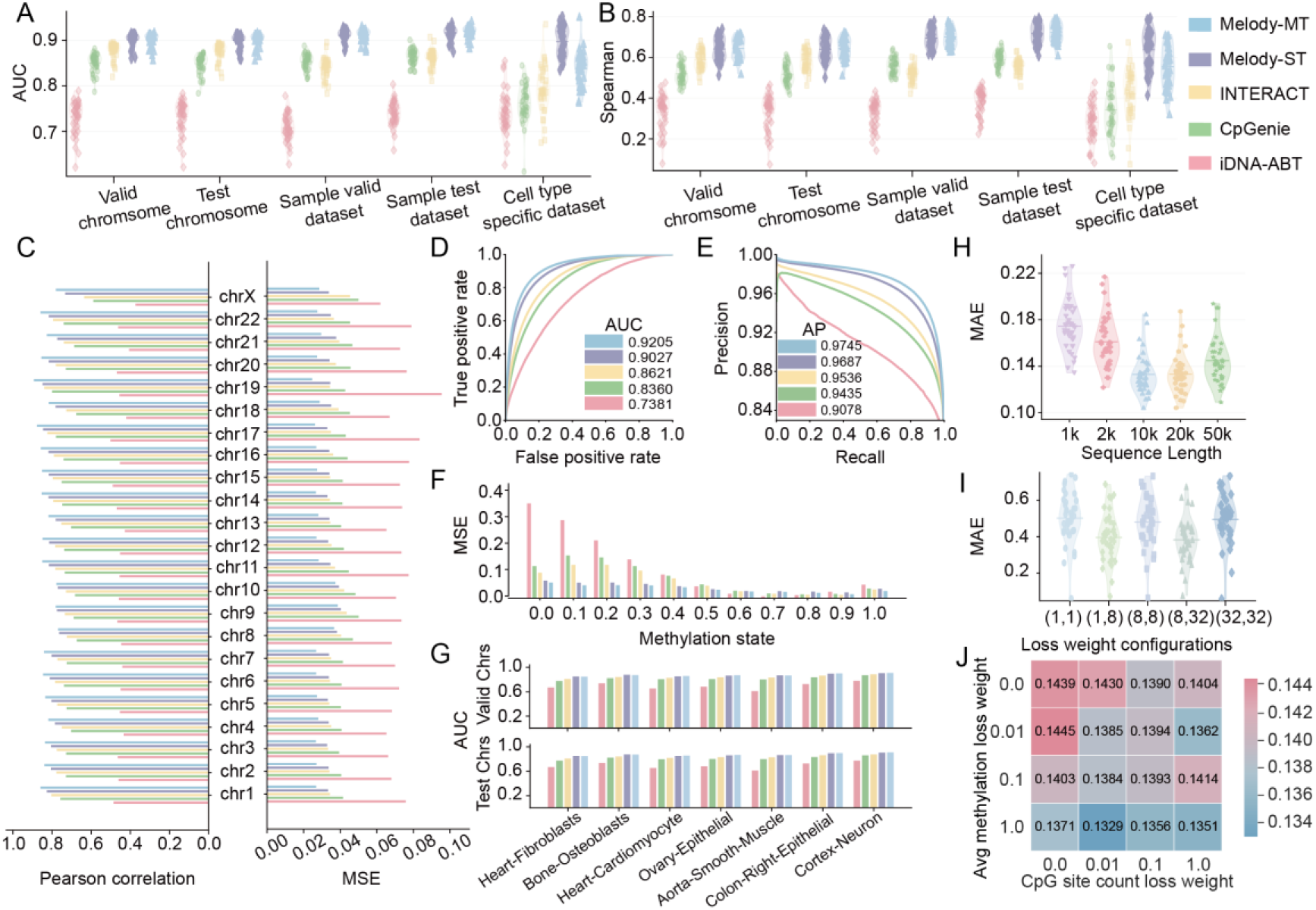
Comparison of model performance with state-of-the-art methods. One point in the figure represents one dataset (n=39). (A–B) Performance comparison across multiple dataset splits, including chromosome- and hypomethylated sample-based valid, test datasets, and cell-type-specific regions. (C) Chromosome level Pearson correlation and MSE comparison across models. (D–E) ROC and precision– recall curves comparison across models. (F) Methylation states level MSE comparison across models. (G) Per-cell AUC on the validation and test chromosomes for seven representative cell types (n=7). (H) Mean absolute error on the validation set across different input sequence lengths. (I) Mean absolute error in cell-type-specific regions across different loss-weight (α, β) configurations. α represents the weight of the CpG site, and β represents the weight of the hypomethylated CpG site. Other non-CpG sites are set to 1. (J) CpG sites counting loss and average methylation level loss weighting comparison quantified by mean absolute error on the validation set averaged across 39 cell types.

The sequence-only model iDNA-ABT, which uses a 41-bp input window, performed substantially worse, indicating that such short sequence contexts are insufficient to capture the broader regulatory information required for accurate methylation prediction. As anticipated, Melody-ST was particularly effective in cell-type-specific regions, surpassing Melody-MT and suggesting an enhanced ability to model localized regulatory effects. Melody-MT achieved the highest area under the ROC curve (AUC, 0.921) and average precision (AP, 0.975) among all methods, with Melody-ST as the second best (AUC 0.903, AP 0.969) (Figure 2D,E). Here, we also present a performance comparison on the validation and test datasets for seven representative cell types, and the results show that Melody consistently outperforms all competing methods (Figure 2G).

To assess robustness, we further evaluated Melody across individual chromosomes and methylation states (Figure 2C,F). On both validation and test chromosomes, Melody attained ∼7% relative improvement in Pearson correlation compared with the best competing model, indicating that its advantages generalize across genomic contexts. When stratified by methylation state, Melody maintained strong Pearson correlations and low mean squared error (MSE) at all methylation levels, demonstrating stable performance. Notably, whereas most baselines deteriorated in hypomethylated regions, Melody remained robust, suggesting that it effectively captures subtle sequence features associated with low methylation levels—regions that are typically difficult to model and biologically complex.

From a model design perspective, we systematically evaluated how performance varies with sequence length, loss-weight configurations, and CpG site count weighting. As shown in Figure 2H and Supplementary Figure S3, the optimal sequence window is around 10 kb, providing sufficient context without excessive parameter overhead; shorter windows (e.g., 2 kb) underperform due to limited contextual information, whereas longer inputs (20–50 kb) introduce redundant sequence regions. Regarding loss weighting, the parameter pair (8, 32)—applied to low-methylation CpG and high-methylation CpG positions, respectively—achieved the lowest MAE in cell-type-specific regions (Figure 2I, Supplementary Figure S4). This setting is consistent with the biological expectation that CpG sites are typically methylated and that the learning objective should prioritize the more challenging low CpG methylation events, with low-methylation CpG positions receiving the highest emphasis. Finally, since our training objective includes both regional average methylation and CpG site count prediction losses, we explored their relative weights and found that assigning higher weight to the regional average methylation loss leads to improved overall performance, while the CpG-count loss provides only a minor additional benefit (Figure 2J, Supplementary Figure S5), consistent with its role as a lightweight structural regularizer. Together, these results demonstrate that Melody effectively captures both local and long-range dependencies, producing accurate, robust, and generalizable DNA methylation predictions across diverse genomic contexts and experimental conditions.

### Melody effectively prioritizes genetic variants influencing DNA methylation

Accurately predicting the impact of genetic variants on DNA methylation is essential for understanding the regulatory mechanisms that connect sequence variation to epigenetic changes. To evaluate this capability, we assessed Melody performance on fine-mapped methylation quantitative trait loci (meQTLs). Because no standardized deep-learning benchmark exists for meQTL prediction, we curated three independent resources, Ólafur et al.^8^, GTEx^7^, and EPIGEN^20^ to construct a more comprehensive and diverse evaluation panel.

We first examined individual meQTL examples using attribution maps^21^ (Figure 3A,B). In Figure 3A, an A→G substitution creates a strong IRF binding motif, which Melody correctly identifies as causing a local decrease in methylation. In Figure 3B, a C→T variant disrupts a CTCF motif, and Melody accurately predicts the resulting increase in methylation. These examples highlight Melody’s ability to infer the direction of methylation change by recognizing allele-specific motif gains and losses. Then for quantitative assessment, we used Pearson correlation between predicted and observed variant effects. Many tissues contain multiple methylation tracks (e.g., GTEx blood may correspond to B cells, T cells, or mixed leukocyte populations), so we first evaluated whether averaging across all related tracks yields more robust prediction than selecting a single track. As shown in Figure 3C, using the average across related tracks (Supplementary Table 2) consistently improves performance for both Melody- MT and Melody-ST. This improvement likely reflects two factors: first, track averaging reduces stochastic noise in the individual methylation maps; second, the averaged signal may better align with the bulk-tissue origin of meQTL evidence, particularly for whole-blood datasets where a single track might represent an overly narrow cell subtype. Given that Melody-ST performs better than Melody-MT on cell-type-specific methylation prediction (Figure 2A–B), we next asked whether this advantage translates to better motif-driven variant effect prediction. Surprisingly, across all meQTLs datasets (Figure 3D), Melody-MT outperforms Melody-ST, suggesting that multi-track architecture provides stronger motif capture ability, whereas ST’s stronger performance in cell-type-specific methylation regions may reflect overfitting to local cell-type patterns rather than learning motif-level regulatory grammar transferable to meQTL prediction.

**Figure 3.**
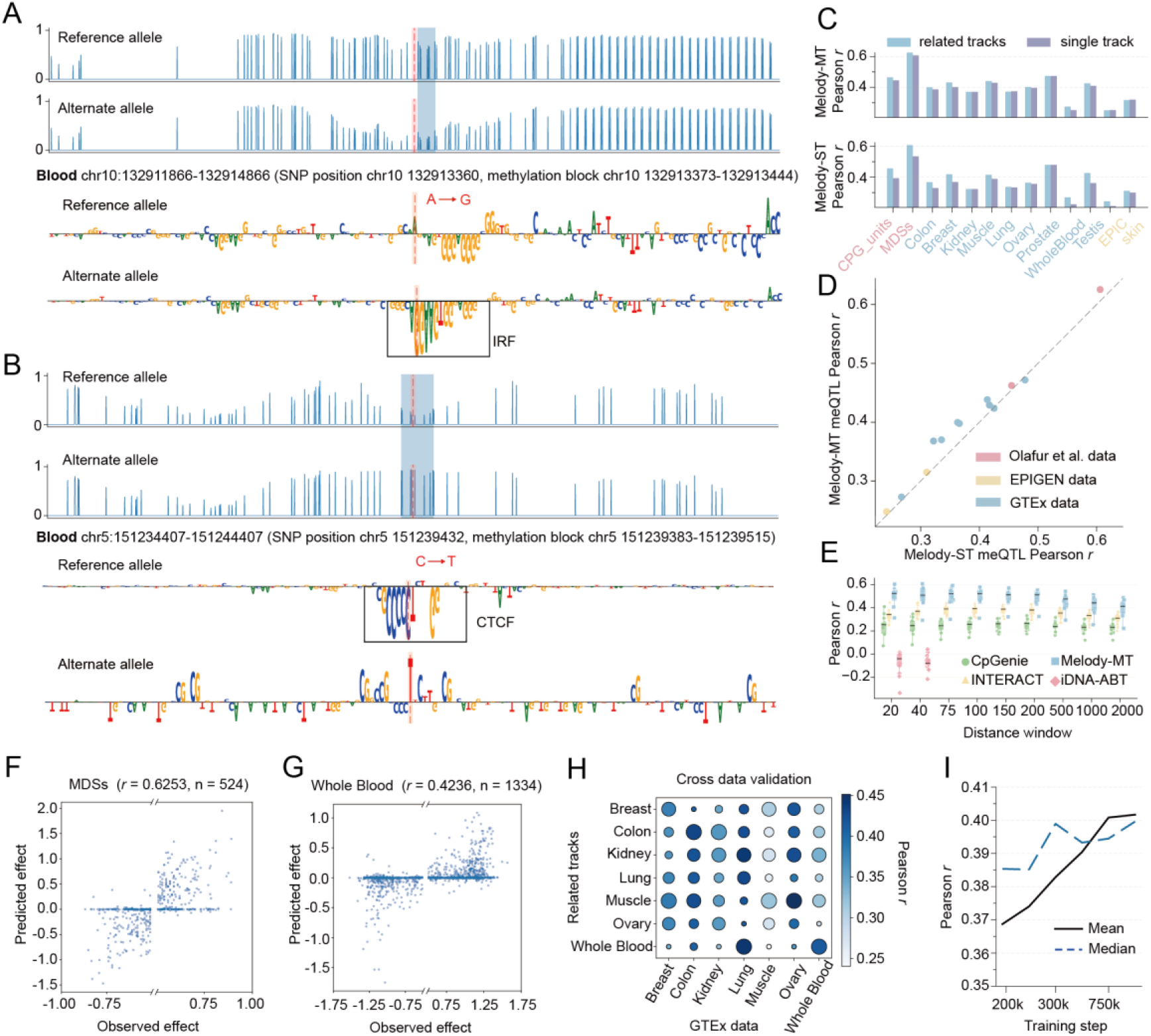
Melody accurately predicts meQTLs effects and recognizes meaningful sequence motifs. (A,B) In silico saturation mutagenesis with Melody-MT for representative meQTLs samples. Predicted methylation profiles are shown before and after the SNP for variants that either increase or decrease methylation levels. The blue- shaded region denotes a cell-type–specific low-methylation zone, and the red dashed line marks the SNP position. (C) Performance of Melody-MT and Melody-ST on meQTLs datasets when using different cell-type–specific prediction tracks. For each dataset, we have two options: either a single representative cell-type channel or the average prediction across several biologically related channels. (D) Comparison of Melody-MT and Melody-ST on meQTLs prediction. (E) Comparison of Melody-MT with other models on all datasets and across different cumulative distance windows. Each point is one of n=13 datasets. The middle black bar indicates the medians; box bounds are the 25th and 75th percentiles; whiskers extend to the minimum and maximum values within 1.5× IQR of the box bounds. (F,G) Scatter plots of Melody-predicted versus observed methylation difference scores for meQTLs in methylation-depleted sequences (MDSs) and the GTEx_WholeBlood dataset.(H) Cross–cell-type evaluation of meQTL prediction on GTEx data. Colour indicates Pearson r, bubble size indicates the rank within each column (largest is best). (I) Relationship between training progress and meQTL prediction performance. Average and median meQTL performance across datasets are shown as a function of training steps.

We then compared Melody to other state-of-the-art models across a range of variants– CpG distances using Pearson correlation (Figure 3E). The direction and absolute magnitude based evaluation of meQTL effect prediction can be found in Supplementary Figure S6. Melody shows superior performance at all distance scales, demonstrating its strong ability to capture meaningful sequence motifs driving methylation variation. In blood datasets such as methylation-depleted sequences (MDSs) and GTEx Blood, Melody achieves Pearson correlations of 0.62 and 0.42, respectively (Figure 3F, G). By contrast, models using short input windows like iDNA-ABT (41 bp) fail to capture meaningful long-range relationships and show near-random performance. All models show decreased performance as variant–CpG distance increases, indicating that long-range regulatory meQTLs are more difficult to explain purely from sequence features—a known limitation given that many distal methylation QTLs involve chromatin contacts and higher-order genomic architecture.

To assess whether Melody’s tissue tracks are biologically meaningful, we conducted cross-track validation (Figure 3H). Related tracks typically achieve the best or second-best performance (e.g., blood, colon), indicating that Melody successfully captures cell-type-specific regulatory motifs relevant to each meQTL dataset. Ovary data perform poorly, likely due to either (i) lower data quality in ovary methylation tracks or (ii) ovary-specific motifs being underrepresented in available meQTL datasets. Finally, we evaluated how training progression affects meQTL prediction (Figure 3I). Both mean and median correlation steadily improve with training steps and converge smoothly, demonstrating that meQTLs prediction is stable throughout training. Models with stronger methylation prediction performance consistently show better meQTL effect prediction, further supporting the link between methylation modeling accuracy and motif capture ability. Together, these results demonstrate that Melody is effective at identifying functional variants that modulate DNA methylation, capturing allele- specific sequence logic with high accuracy across tissues, datasets, and genomic distances.

### Melody reveals motif variability and regulatory differences across tissues

To dissect how sequence-resolved perturbations influence DNA methylation across regulatory contexts, we performed a motif-centered allelic perturbation analysis using Melody (Figure 4A). To comprehensively evaluate motif effects, we curated 282 transcription factor motifs from a motif database^22^. For each motif, we randomly sampled one representative instance from the database. To ensure meaningful perturbation signals, we sampled genomic methylation blocks containing at least four CpG sites from the training dataset. Motifs were then inserted into regions spanning 600 bp upstream of the block start and 600 bp downstream of the block end. The difference between the predicted methylation level of the motif-inserted sequence and the original unmodified sequence was defined as the motif effect at that position. We used Melody-MT as the primary predictor for this analysis because it demonstrated the strongest performance in our meQTLs benchmark.

**Figure 4.**
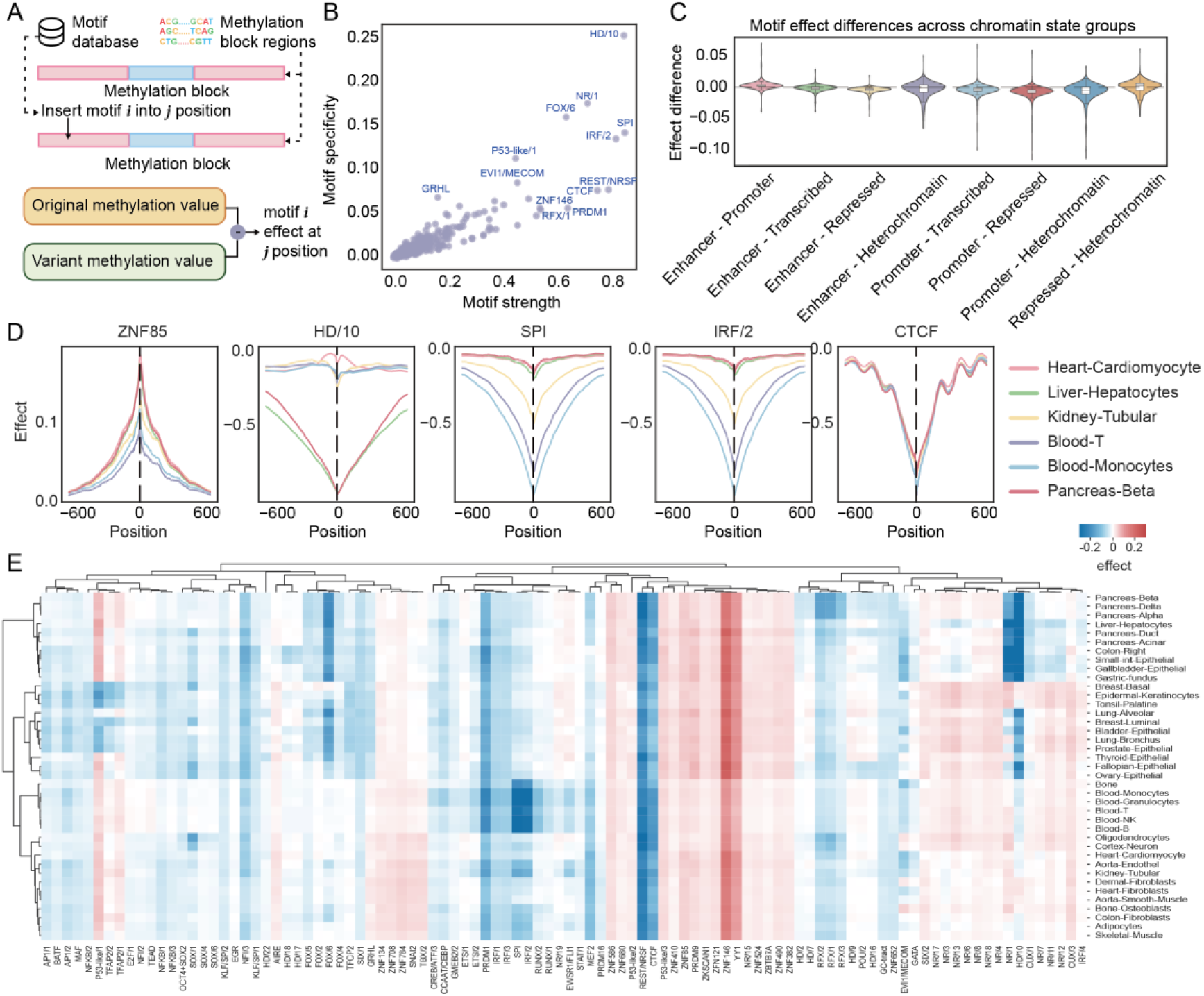
Melody maps motif-driven methylation variation across tissues. (A) Overview of the motif-centered allelic perturbation workflow comparing methylation predictions between reference and motif-inserted sequences. (B) Scatterplot showing motif strength and tissue specificity across 282 transcription factor motifs. (C) Violin plot displaying the distribution of 282 transcription factor motif methylation effects across chromatin states. Each value is the difference, for one motif, between its mean motif effect in the two chromatin-state groups named on the x-axis. The middle line indicates the median value, box bounds indicate Q1 and Q3, whiskers indicate most extreme point within 1.5 x IQR, outliers not drawn. (D) Positional effect profiles for representative motifs (ZNF85, HD/10, SPI, IRF/2, CTCF) across six tissues. (E) Heatmap showing methylation effects across different motifs and tissues.

By predicting methylation landscapes for both reference and variant sequences, Melody isolates the contribution of individual motifs and quantifies their allele-specific positional effects with high resolution. To be noted, these results highlight the associations between motifs and methylation patterns rather than establishing definitive causal relationships. Several example motifs are shown in Figure 4D. Across six representative tissues selected from our 39-tissue panel, motifs exhibited distinct effect profiles: ZNF85 produced a positive local methylation shift, whereas HD/10, SPI, IRF2, and CTCF produced negative shifts. Notably, tissue-specific differences are also apparent—HD/10 exhibits stronger effects in liver and pancreas, while SPI and IRF2 show their strongest signals in blood-derived cell types. The CTCF motif displays a particularly characteristic non-smooth, oscillatory pattern reminiscent of nucleosome phasing, consistent with reported CTCF–nucleosome interactions. Melody predicts a pronounced local loss of methylation proximal to the CTCF motif, aligning with CTCF’s known function as a methylation-sensitive insulator that protects CpGs from DNMT activity and promotes demethylation via TET recruitment^23^.

To better quantify motif differences, we defined two metrics: strength, measuring how strongly a motif perturbs methylation (computed as the maximal absolute effect in the positional curve), and specificity, measuring cell-type variation (computed as the variance of effects across tissues). As shown in Figure 4B, motifs such as HD/10 and NR/1 exhibit both high strength and high tissue specificity, highlighting their potential roles as lineage-restricted regulators of methylation landscapes. Previous studies suggest that methylation responsiveness varies substantially across chromatin states, particularly in promoter and enhancer regions, our motif-level analysis revealed more nuanced trends. When motif effects were aggregated by consolidated chromatin states (Promoter, Enhancer, Transcribed, Repressed, Heterochromatin; Figure 4C), promoter regions indeed showed the largest magnitude of variation. However, enhancer regions were less distinct than often assumed, and repressed regions displayed unexpectedly strong and coherent effects that were markedly different from enhancers—suggesting fundamentally different regulatory mechanisms. Despite these differences, motif-driven methylation responses remained broadly stable across chromatin states, indicating that sequence-intrinsic biochemical interactions between transcription factor motifs and the methylation machinery dominate over chromatin context.

Finally, to evaluate how well Melody captures cell-type-specific regulatory logic, we generated a motif-by-cell-type heatmap (Figure 4E). This analysis revealed clear cross-tissue perturbation modules: global architectural regulators such as CTCF and REST formed broad, pan-tissue clusters, whereas blood-derived cell types clustered strongly and were dominated by hematopoietic lineage factors SPI and IRF2. These patterns recapitulate known transcriptional hierarchies and demonstrate that Melody effectively captures both global and lineage-specific regulators of DNA methylation. Collectively, these results show that Melody provides an interpretable, sequence-resolved framework for prioritizing genetic variants that modulate DNA methylation across diverse chromatin and cellular contexts.

### Melody can generalize to unseen tissues and accurately captures cell type specific methylated regions

Unlike Melody-MT and Melody-ST, to evaluate Melody-G’s generalization performance, we designated five cell types (Aorta-Endothel, Blood-Granulocytes, Blood-Monocytes, Cortex-Neuron, and Pancreas-Delta) as an unseen test set, training the model on the remaining 34 cell types. As a baseline, we used the average methylation profile of the 34 seen cell types, computed by averaging methylation probabilities across these cell types at each CpG site. On the unseen set, Melody-G was highly effective (Figure 5A). Its two training stages, G1 and G2, achieved mean AUCs of 0.663 and 0.697, respectively. These values represent improvements of 4.7% and 10.1% over the MT- mean baseline (AUC = 0.633). This result confirms that integrating scRNA-seq embeddings provides strong guidance for predicting cell-type-specific methylation patterns. Conversely, when evaluated on the seen cell types (Figure 5B), both G1 and G2 showed a slight performance degradation relative to single cell type models. This is an expected trade-off: whereas models like Melody-ST and Melody-MT are specialized for a single cell type, Melody-G performs a more complex multi-task objective of concurrently inferring cell identity and predicting methylation. This powerful generalization capability necessarily entails a modest performance cost on the seen cell types. Furthermore, because G2 was fine-tuned exclusively on cell-type-specific regions, the model’s focus shifted toward localized regulatory signatures. As a result, its capacity to model broad, non-specific background patterns was comparatively diminished, leading to the slight performance decrease observed in this evaluation setting. Consistent with this specialization trade-off, G1 maintained overall superiority over INTERACT, whereas G2’s global performance was slightly weaker (Figure 5B).

**Figure 5.**
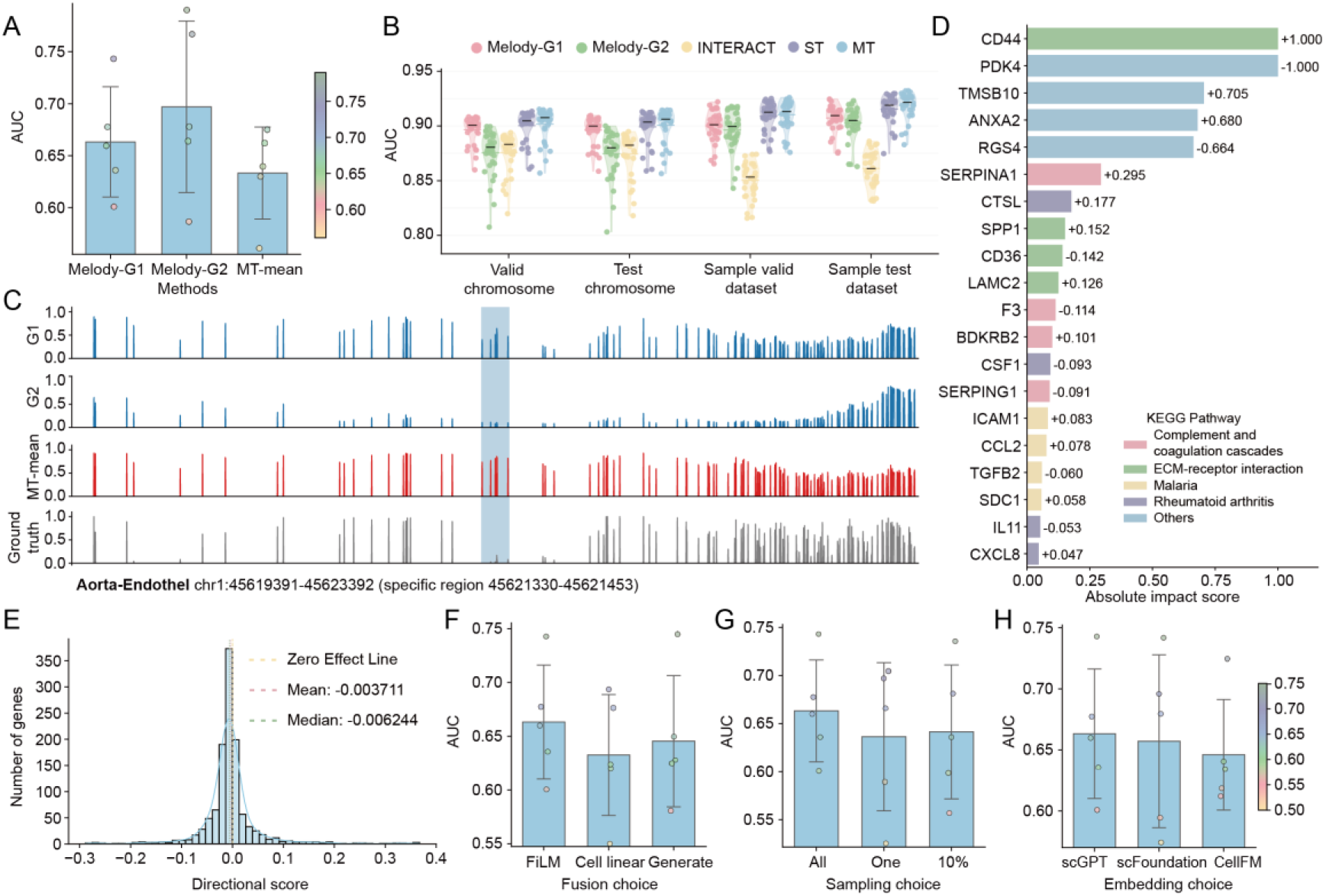
Performance analysis of Melody-G driven by scRNA-seq embeddings. (A) Performance comparison of Melody-G’s two training stages (G1, G2) versus the mean-based MT model (MT-mean). Scatter points represent the performance distribution across n=5 unseen cell types. Bars show the mean AUC across five unseen cell types; error bars indicate the standard deviation; points represent individual unseen cell types. (B) Performance comparison across different methods on seen cell types (n=34) under chromosome-based, sample-based validation, and test-set evaluations. Violin plots show the distribution of AUCs across seen cell-type tracks; dots represent individual tracks, colored horizontal lines indicate the mean, and black ticks indicate the median. (C) Predicted methylation profiles from Melody-G (blue) and the mean-normalized MT model (red) are compared with the ground truth profile (gray). The profiles span a cell-type-specific region (highlighted by blue shading) and its 2000 bp flanking regions. (D) Functional annotation of genes ranked by influence score. Genes (y-axis) are sorted by the absolute value of their directional influence score, with the magnitude indicated by bar length (x-axis) and the direction retained in the signed bar-end labels. Bars are color-coded according to their membership in the four representative enriched KEGG pathways. Genes not associated with these pathways are shown in blue (’Others’). (E) Histogram with a kernel density estimate (KDE) of directional influence scores for highly variable genes (HVGs). The influence score (x- axis) is defined as the change in the model-predicted mean CpG methylation upon in silico gene knockout (mean-knockout – mean-baseline). The y-axis represents the number of genes. For visualization, the central 98% of the score distribution is shown. Vertical dashed lines indicate the distribution’s mean (purple), median (green), and a zero-effect reference (yellow). (F) Performance comparison of different scRNA-seq embedding fusion strategies. (G) The effect of different scRNA-seq sample sizes on model performance. (H) The effect of embeddings generated by different scRNA-seq foundation models on model performance. For (F-H), scatter points represent the performance distribution across n=5 unseen cell types. Bars show the mean AUC across five unseen cell types; error bars indicate the standard deviation; points represent individual unseen cell types.

To visualize performance in unseen cell-type-specific regions, we examined the predicted methylation profile for a representative locus in Aorta-Endothel (Figure 5C). Melody-G2 accurately captured the characteristic hypomethylation pattern within the cell-type-specific region, demonstrating its specialized learning. However, its accuracy was diminished in the flanking non-specific regions. This suggests that while G2’s fine-tuning enhances specificity, it comes at the cost of global predictive accuracy. In contrast, Melody-G1 demonstrated robust and balanced performance across both specific and non-specific regions, making it the more suitable model for generating complete and reliable methylation profiles in unseen cell types.

To interpret the gene-level drivers of our model’s predictions, we performed in silico gene knockouts using Pancreas-Delta scRNA-seq data to quantify each gene’s influence. Subsequent KEGG analysis of representative genes revealed enrichment in distinct biological pathways (Figure 5D)^24,25^. For example, CTSL and CSF1 were selected as representative model-responsive genes annotated to the rheumatoid arthritis pathway; CTSL has been implicated in rheumatoid synovial fibroblast– mediated cartilage destruction and macrophage-associated subchondral bone erosion, whereas CSF1–CSF1R signaling promotes osteoclast differentiation and contributes to bone erosion in experimental arthritis^26–29^. This finding suggests that the scRNA-seq embeddings capture the functional state of critical pathways, providing a mechanistic basis for the model’s predictive efficacy.

We next investigated how these knockouts affected the predicted methylation levels (Figure 5E). Across the 1,200 representative genes, changes in mean predicted CpG methylation were bidirectional: approximately two-thirds of the gene perturbations decreased the model-predicted mean CpG methylation relative to the unperturbed baseline, whereas approximately one-third increased it. These results indicate that Melody-G1 responds to transcriptomic features in a gene-specific and direction-dependent manner, providing a model-level view of how the scRNA-seq embedding contributes to methylation prediction.

To dissect the contributions of our scRNA-seq embedding strategy, we conducted a series of ablation studies (Figure 5F-H and Supplementary Figure S7). First, we compared different methods for integrating the embeddings and found that feature-wise linear modulation (FiLM) showed the best overall performance in our benchmark, although its relative advantage depended on the upstream foundation model and the evaluation split. It achieved an AUC of 0.663, outperforming both a simple linear embedding and a parameter generation approach by 4.7% and 2.8%, respectively (Figure 5F). We also confirmed that comprehensive cellular data is critical, as using randomly sampled subsets of cells led to a marked decrease in performance (Figure 5G). Furthermore, a full cross-ablation across foundation models, fusion strategies, and embedding sampling settings (Supplementary Figure S7A) revealed that FiLM remained the optimal fusion strategy; however, while strong and consistent gains were observed for scGPT and scFoundation, the relative advantage was notably weaker for CellFM.

To further disentangle the contribution of RNA embeddings from that of sequence window length, we additionally evaluated Melody-G under shorter input windows of 1 kb and 2 kb, matching the scale of previous sequence-only models. Under both matched window settings, Melody-G consistently outperformed the corresponding Melody-MT baseline (Supplementary Figure S7B). This indicates that the predictive gain is not solely driven by the extended 10 kb sequence window, but fundamentally reflects the added value of RNA embeddings from single-cell foundation models. Finally, when comparing the three scRNA-seq foundation models, scGPT showed the strongest overall performance in the original benchmark (Figure 5H). However, their relative ranking was not fixed across evaluation settings. In particular, under a more stringent unseen split composed of transcriptomically distinct cell types (Supplementary Figure S8), scFoundation+FiLM achieved the best performance (Supplementary Figure S7C). Together, these results demonstrate that the utility of the upstream cell representation heavily depends on both the held-out test scenario and the degree of transcriptomic shift between the training and test sets.

## Discussion

In this work, we present Melody, a deep learning framework for modeling DNA methylation directly from genomic sequence across diverse cellular contexts. By combining base-resolution prediction, multi-track learning, and transfer to unseen cell types through cell-state embeddings derived from single-cell foundation models, Melody provides a unified framework for sequence-based methylation prediction and downstream variant-effect analysis. Our results show that incorporating broader sequence context and cell-type-aware information improves predictive performance across tissues and cell types, supporting the view that a substantial component of DNA methylation patterning is encoded in the underlying DNA sequence.

Beyond predictive accuracy, Melody provides a framework for interrogating the sequence features associated with methylation variation and for evaluating the impact of genetic variation on methylation states. In this sense, the model serves not only as a predictor, but also as a computational tool for studying the sequence basis of epigenomic variation. These findings add to growing evidence that deep learning models can recover meaningful regulatory signals from DNA sequence alone, while also highlighting the extent to which methylation regulation reflects both shared and cell-type-specific sequence logic.

Several limitations should be noted. First, although DNA sequence provides an important substrate for methylation regulation, methylation states are also shaped by chromatin accessibility, transcription factor occupancy, developmental history, replication timing, and other aspects of cellular context. As a result, a sequence-only framework cannot fully capture all sources of methylation variability, particularly in highly context-dependent settings. Second, the interpretability analyses presented here should be understood as model-based assessments of learned sequence associations rather than direct evidence of causal regulatory mechanisms *in vivo*. Third, generalization to unseen cell types remains constrained by the quality, coverage, and granularity of currently available reference methylation and cell-state datasets.

These limitations point to several important opportunities for future development. One natural direction is to incorporate broader biological context into methylation modeling, for example by integrating additional epigenomic signals such as chromatin accessibility, histone modifications, or transcription initiation. Such extensions may help clarify how DNA methylation relates to other layers of regulatory activity and may improve prediction in contexts where sequence alone is insufficient. More generally, an important next step will be to move beyond association-based modeling toward a more explicit understanding of the causal relationships among DNA sequence, DNA methylation, chromatin accessibility, and transcriptional regulation. While sequence-based models can identify features associated with methylation variation, they do not by themselves distinguish whether methylation is a driver, a consequence, or a correlated marker of regulatory activity. Joint modeling of multiple molecular layers may therefore help disentangle these relationships and provide a more mechanistic view of how sequence variation propagates through epigenomic state to transcriptional output.

A second direction is methodological: larger-scale pretraining strategies, including those informed by evolutionary conservation or other genome-wide regulatory priors, may further improve representation learning and adaptation to methylation-related tasks. More generally, it will be of interest to explore whether newer model backbones or pretrained genomic foundation models can be effectively adapted to methylation prediction through task-specific fine-tuning.

Finally, the broader implications of such models may extend beyond basic epigenome prediction. Because DNA methylation is widely used as a biomarker of aging, disease state, and cellular identity, improved sequence-based models may provide a useful foundation for studying disease-associated methylation variation, including cancer-related epigenetic alterations and pathogenic meQTLs effects. Although substantial work remains to establish translational utility, frameworks such as Melody may help bridge sequence-based regulatory modeling with the study of human disease.

Overall, Melody provides a general and interpretable framework for modeling DNA methylation from sequence across cellular contexts. Our study highlights both the promise and the current limitations of sequence-based methylation modeling, and establishes a foundation for future work linking DNA sequence, epigenomic state, genetic variation, and cellular identity in a unified predictive framework.

## Methods

### Dataset

Training and evaluation datasets were derived from the DNA methylation atlas of normal human cell types^18^. To construct a representative and diverse training set, we selected 39 distinct normal human cell types from this atlas (Supplementary Table 1), prioritizing samples with clear lineage and functional representation. All data underwent standard quality control procedures to ensure the consistency of methylation measurements. Melody takes genome sequences, like reference genome (FASTA, e.g., GRCh38 primary assembly), as input and predicts base-resolution methylation tracks in BigWig format aligned to the same assembly; scRNA-seq input for Melody-G is provided as a cell × gene expression matrix (AnnData/h5ad). Data input and model training was implemented using Selene^30^.

### Model

The core architecture of Melody is based on a 1D U-Net topology^31^, which has proven effective for modeling biological sequence data with hierarchical spatial structure in prior work such as AlphaGenome and Enformer. The U-Net model consists of a symmetric encoder–decoder with successive downsampling and upsampling blocks connected by skip connections, allowing the network to integrate long-range context while preserving base-pair–resolution local information. Here, to better adapt the U-Net architecture to the DNA methylation prediction task, we modify several architectural components, including the normalization layers and convolutional kernels. The model takes base-pair–resolution nucleotide sequences as input and uses a sigmoid output layer to produce base-pair–resolution DNA methylation probabilities. To enhance the model’s ability to capture complex DNA methylation patterns, we introduced several key methodological innovations:

Auxiliary Loss Functions. To ensure the model learned relevant genomic features at multiple scales, we incorporated two auxiliary loss terms alongside the primary prediction task. The first, termed average loss, compels the model to predict the mean methylation level within each 100 bp segment. This loss is as a supplement for region methylation capture. The second, CG loss, tasks the model with predicting the total count of CpG dinucleotides (i.e., CpG density) within the same segment. This loss works as a CpG island capture. This multi-task learning strategy was found to significantly stabilize training and improve overall model performance.

Weighted Loss Mechanism. Recognizing the functional heterogeneity of different CpG sites, we implemented a weighted loss function. This function is governed by two hyperparameters, α and β, which selectively increase the loss contribution of all CpG sites (α) and, more specifically, low-methylation CpG sites (β). By focusing the model’s attention on these sites, which are often enriched in active regulatory elements (e.g., promoters and enhancers), this weighting scheme demonstrably improved predictive accuracy and enhanced the model’s sensitivity in downstream meQTL analyses.

We developed three configurations of the Melody framework to address distinct prediction scenarios: Melody-ST (single-track), Melody-MT (multi-track) and Melody-G (generalize). All three share the same sequence-to-methylation backbone described above, but differ in how they use cell-type information. Melody-ST and Melody-MT operate on cell types with directly measured methylation tracks, specializing either in a single cell type or in joint multi-cell-type modeling. Melody-G extends this design by additionally conditioning on cell embeddings derived from scRNA-seq data, enabling methylation prediction in unseen cell types.

### Model Configuration and Training

Melody was trained on random genomic windows sampled from the GRCh38 primary assembly using a custom sampler built on the Selene framework. At each training step, we drew a 10-kb window (10,000 bp) from a random genomic position and retrieved the one-hot–encoded nucleotide sequence together with the corresponding base-resolution methylation tracks for all available cell types from bigWig files. The primary prediction loss was defined at base resolution and computed over all valid positions in each batch. All models were trained with a batch size of 32 using the Adam optimizer on a single NVIDIA A100 GPU (80 GB); a typical training run completed in approximately 72 hours.

### Melody-ST

Melody-ST is a single-track configuration tailored for cell-type-specific methylation prediction. For a given experiment, the model is trained on one cell type at a time, using a single bigWig track as supervision, and produces a base-resolution methylation probability for each genomic position. Architecturally, Melody-ST uses the 1D U-Net backbone with a single output channel (track = 1) in the final 1×1 convolution layer, and retaining the auxiliary heads that predict CpG counts and regional methylation levels at 100 bp resolution.

### Melody-MT

Melody-MT is a multi-track configuration designed to jointly model DNA methylation across multiple cell types. In this setting, the model is trained with 39 methylation tracks from the atlas, and the final 1×1 convolution layer outputs track = 39 channels, each corresponding to one cell type. All cell types share the same encoder–decoder parameters, while their methylation profiles are disentangled in the channel dimension of the output. The auxiliary 100 bp heads, predicting CpG density and average methylation, are defined over the same shared feature maps and output one value per segment for each track, providing additional supervision on both CpG architecture and regional methylation patterns. This multi-task training encourages the backbone to learn regulatory features that are partially shared across tissues while still allowing track-specific specialization. In practice, Melody-MT serves as the default model for tasks that involve multiple tissues, including meQTLs effect prediction and cross–cell-type transfer, and also provides the mean-track baseline (MT-mean) used to evaluate Melody-G on unseen cell types.

### Melody-G

To enable the prediction of DNA methylation in unseen cell types, our model integrates DNA sequences with cell-type-specific features derived from scRNA-seq data. This approach provides the necessary cellular context for the model to generalize and make cell-type-specific predictions, even for cell types not encountered during training.

### The scRNA-seq Data Collection and Preprocessing

Single-cell RNA-seq datasets for 39 cell types were obtained from the National Center for Biotechnology Information (NCBI). All relevant accession numbers are listed in Supplementary Table 3. Raw scRNA-seq gene expression counts were processed using the Scanpy (v1.11.1)^32^ package. The preprocessing workflow involved two main quality control steps: first, we filtered the dataset to a predefined panel of 19,264 target genes by adopting the strategy of Hao et al^33^. Second, we removed cells expressing fewer than 200 genes. The resulting quality-controlled gene-cell count matrix served as the input for all downstream analyses.

### Generation of cell embeddings

We utilized three scRNA-seq foundation models to generate cell embeddings from gene expression data: scGPT^34^, scFoundation^33^, and CellFM^35^. For scGPT, we generated cell embeddings using the pre-trained scGPT model, following the methodology of Cui et al. First, we selected the top 1,200 highly variable genes (HVGs) and discretized their log-normalized expression values into 51 bins. We then converted each cell’s profile into a sequence of tokens. A special <cls> classification token was prepended to each sequence to enable the model to learn a global cell representation. After processing these sequences with the scGPT model, we extracted the output embedding corresponding to the <cls> token for each cell. Finally, these raw embeddings were L2-normalized to produce the final cell representations used in all downstream analyses.

For scFoundation, we generated cell embeddings using the pre-trained scFoundation model, following the methodology of Hao et al^33^., from a quality-controlled expression matrix of 19,264 genes. For each cell, the model first constructs an input representation for each gene by summing two distinct components. The first component is a gene name embedding retrieved from a learnable lookup table, while the second is an expression value embedding derived from its continuous, non-discretized value. A Transformer architecture then processes the full sequence of these combined gene representations to produce a single, global embedding for each cell. These final embeddings served as the input for all downstream analyses.

For CellFM, we generated cell embeddings using the pre-trained CellFM model, following the methodology of Zeng et al^35^. A key preprocessing step involved standardizing each cell’s profile to a fixed-length sequence of 2,048 genes. This was achieved by selecting the 2,048 most highly expressed genes for cells exceeding this limit and by padding with zero-value tokens for cells below it. The model then constructs an input representation for each gene by summing two components: a value embedding derived from its continuous expression value via a multi-layer perceptron, and a gene identity embedding retrieved from a learnable lookup table. The resulting sequence of combined embeddings was then processed by the model’s core architecture to generate the final cell embeddings.

### Data preparation for training

For each cell type, we generated a representative embedding by averaging the individual embedding vectors from all of its constituent cells. To assess the impact of data scale on performance, we then evaluated two alternative sampling strategies. The first setting used an embedding from a single, randomly selected cell. The second used the average of embeddings from a randomly sampled 10% of the cell population.

### Two-stage training strategy

We designed a two-stage training protocol for Melody-G to evaluate its cross-cell-type generalization. First, we partitioned the 39 available cell types into a training set of 34 “seen” types and a held-out test set of five “unseen” types (Aorta-Endothel, Blood- Granulocytes, Blood-Monocytes, Cortex-Neuron, and Pancreas-Delta). The first training stage (G1) aimed to learn general genomic patterns from the seen cell types, using chr10 for validation, chr8 and chr9 for testing, and the remaining chromosomes for training. The second stage (G2) then focused on fine-tuning for cell-type specificity by training exclusively on cell-type-specific regions from the seen types and evaluating performance on the corresponding regions in the five unseen cell types.

### Cell-conditioned feature fusion

To integrate DNA sequences with scRNA-seq data, we evaluated three distinct strategies: feature-wise linear modulation (FiLM)^36^, cell linear addition, and a generated linear layer parameters approach.

Feature-wise Linear Modulation (FiLM) Layer. To integrate cell-type-specific contextual information into our model’s feature processing pipeline, we implemented a Feature-wise Linear Modulation (FiLM) layer. This module conditions the intermediate feature representations on a sample-specific basis. Let *x* ∈ *R^C^*^×*L*^ represent an intermediate feature tensor, with *C* channels and a sequence length of *L*. Let *e_cell_* ∈ *R^dcell^* be the corresponding cell embedding vector, where *d_cell_* is the dimension of the embedding. The purpose of the FiLM layer is to dynamically generate channel-wise affine transformation parameters from *e_cell_* and apply them to *x*.

The modulation parameters are generated by a dedicated two-layer Multi-Layer Perceptron (MLP) which takes the cell embedding *e_cell_* as input. The first hidden layer of the MLP projects the embedding into a space of dimension 2*C* and applies a Rectified Linear Unit (ReLU) activation. This is followed by a second linear layer that maps this intermediate representation to the final conditioning vector. This process can be formally described as:

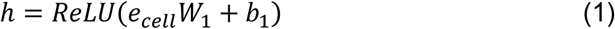

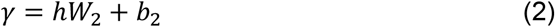

where *W*_1_ ∈ *R^dcell^*^×2*C*^, *b*_1_ ∈ *R*^2*C*^, *W*_2_ ∈ *R*^2*C*×2*C*^, and *b*_2_ ∈ *R*^2*C*^ are learnable weight matrices and bias vectors of the MLP. The resulting output, *γ* ∈ *R*^2*C*^, contains the concatenated modulation parameters for all channels.

The conditioning vector *γ* is subsequently partitioned into two distinct vectors: a channel-wise scaling factor, *s* ∈ *R^C^*, and a shifting factor, *β* ∈ *R^C^*, such that *γ* = [*s*′ ∥ *β*′], where ∥ denotes concatenation. These parameters are then used to perform a channel-wise affine transformation on the input tensor *x*. To enable broadcasting across the length dimension *L*, *s* and *β* are reshaped to *s*′ ∈ *R^C^*^×1^ and *β*′ ∈ *R^C^*^×1^, respectively. The final modulated feature tensor, *x_modulated_*, is computed as:

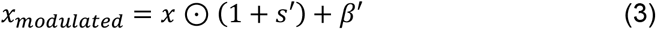

where ⊙ denotes the element-wise Hadamard product.

Cell linear addition. To incorporate cellular context into our model, we employed a direct feature fusion mechanism. An intermediate feature tensor, *x* ∈ *R^C^*^×*L*^, was first obtained from the output of a convolutional layer. In parallel, the original cell embedding vector, *e_cell_* ∈ *R^dcell^*, was processed by a linear layer to project it to the same channel dimension *C* as the feature tensor. This projection can be described as:

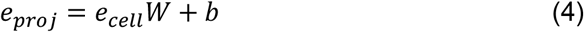

where *W* ∈ *R^dcell^*^×*C*^ and *b* ∈ *R^C^* are learnable parameters.

The resulting projected vector, *e_proj_* ∈ *R^C^,* was then fused with the feature matrix. To achieve this, the vector is broadcast across the length dimension *L* and added element-wise to *x*. This operation can be explicitly written by first reshaping the vector to *e*′*_proj_* ∈ *R^C^*^×1^:

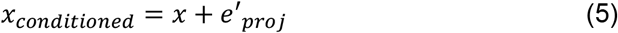

Generated linear layer parameters. To achieve a highly adaptive, cell-type-specific transformation, we employed a hypernetwork architecture. This approach dynamically generates the parameters (weights and biases) of a linear layer based on the cellular context provided by a cell embedding vector.

Let *x* ∈ *R^C^*^×*L*^ be an intermediate feature matrix for a single sample, with *C* channels, and let *e_cell_* ∈ *R^dcell^* be the corresponding cell embedding, where *d_cell_* is the embedding dimension. A parameter generation network, denoted as a function *f_θ_*, maps the cell embedding to a flattened vector containing all parameters required for a target linear layer projecting from *C* input features to *C_out_* output features. The generator *f_θ_* is a multilayer perceptron (MLP) with GELU activations. The generation process is given by:

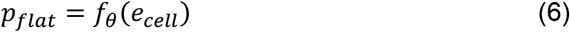

where the output *p_flat_* ∈ *R^Cout^*^×(*C*+1)^ contains the concatenated weight matrix and bias vector.

This flat parameter vector is then reshaped and partitioned to yield the dynamic weight matrix *W_dyn_* ∈ *R^Cout^*^×*C*^ and the dynamic bias vector *b_dyn_* ∈ *R^Cout^* . The dynamically generated linear layer is then applied to the input feature matrix *x.* This is achieved via a matrix multiplication between the dynamic weights and the input features, followed by the addition of the dynamic bias:

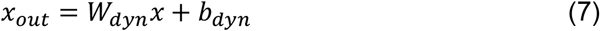

### Baseline settings

To benchmark our approach, we reproduced three representative methylation prediction methods, specifically CpGenie^14^, iDNA-ABT^15^, and INTERACT^17^. Given that these are site-level models, we extracted chromosomal CpG sites for training and evaluation to ensure a fair comparison while strictly adhering to the original model architectures. CpGenie utilizes a CNN composed of three convolutional layers with 128, 256, and 512 channels followed by max-pooling to extract features from 1001 bp input sequences. iDNA-ABT employs a 3-layer BERT architecture with a hidden dimension of 32 designed for 41 bp sequences. Finally, INTERACT processes 2 kb DNA sequences by first extracting features via a 512-channel 1D-CNN coupled with max-pooling, and then feeding these embeddings into an 8-layer BERT encoder to predict the final methylation level through a fully connected layer. We did not benchmark iDNA-ABF^16^ because its pretrained weights exhibited numerical instability during fine-tuning. Since iDNA-ABF shares the same 41-bp input scale as iDNA-ABT, iDNA-ABT serves as a representative baseline for that class of short-window models.

### Performance comparison

Melody variants were optimized using a chromosome-wise data split wherein chr10 served as the validation set, chr8 and chr9 constituted the test set, and the remaining chromosomes were utilized for training across their respective cell types. To better evaluate the model performance, we add two small datasets, one is sampling from hypomethylated regions called sampling dataset, the other is sampling from cell-type-specific datasets. The training procedure for Melody-G adhered to the aforementioned two-stage training strategy. For performance assessment, we focused exclusively on CpG sites, quantifying predictive capability via AUC and Accuracy metrics within the designated evaluation regions.

### meQTLs datasets

We assembled a unified multi-cohort resource by integrating three independent datasets: Ólafur et al., GTEx, and EPIGEN. From the Ólafur et al. study^8^, we obtained fine-mapped CpG-level meQTLs and coordinates of methylation-depleted sequences (MDSs). The GTEx dataset^7^ comprised cis-meQTLs mapped across nine tissues (breast, colon, kidney, lung, skeletal muscle, ovary, prostate, testis, whole blood) from 987 individuals. The EPIGEN resource (https://epicmeqtl.kcl.ac.uk/) included two complementary databases curated by the Epigenomics Research Group at King’s College London: (i) the EPIC database^20^, reporting meQTLs at 724,499 CpGs in 2,358 blood samples profiled using the Illumina EPIC array, and (ii) the Skin database, containing conditionally independent meQTLs identified in whole-skin tissue from up to 394 twins profiled with the Illumina 450K array.

All datasets were harmonized to GRCh38 and standardized into a common variant– CpG format. To ensure high-confidence associations across platforms and cohorts, we retained only variant–CpG pairs with p < 1 × 10⁻⁵ and an absolute methylation effect size greater than 0.5. These filters reduce spurious associations and emphasize variants with reproducible, biologically meaningful methylation changes. The resulting benchmark spans diverse tissues, methylation platforms, and linkage structures, providing a comprehensive and stringent evaluation set for assessing deep-learning models on meQTL effect prediction.

### Motif-centered allelic perturbation analysis

Transcription factor motifs were obtained from the comprehensive compendium and grouped according to their archetype clusters^22^. For each cluster, we extracted the consensus sequence and converted it into a 4×L one-hot representation to enable precise sequence-level perturbations in the model input.

To assess how individual regulatory motifs shape local DNA methylation patterns, we performed a systematic motif-insertion perturbation analysis across all selected CpG-containing regions. For each locus, a 10-kb window centred on the region was used to obtain baseline predictions from the multi-track methylation model, providing unmodified methylation probability profiles across 39 tissues. Motifs were then inserted into the same sequence at regularly spaced positions extending ±600 bp from the CpG interval, using 5-bp increments to capture fine-scale positional sensitivity. Insertions were executed both upstream and downstream of the CpG window, with genomic coordinates adjusted individually for each region.

Following insertion, all modified sequences were re-evaluated with the model under identical conditions, generating perturbed methylation predictions for every tissue. The methylation effect of each motif was computed as the change in predicted methylation within the CpG interval relative to the unmodified sequence. Because CpG density varies across regions, all perturbation effects were normalized by the number of CpGs within each locus, yielding a position-specific, tissue-resolved estimate of the methylation shift attributable to motif insertion. This framework enabled a high-resolution quantification of motif influence on DNA methylation across diverse regulatory contexts.

### KEGG

Quantifying the impact of each gene on methylation prediction involved an in silico perturbation approach. A baseline prediction was first generated using the original, unperturbed cell embeddings. Next, we systematically evaluated pre-computed embeddings corresponding to single-gene in silico knockouts. A directional impact score was then calculated for each gene as the difference between the mean prediction in the perturbed state and the baseline. A positive score indicated an inhibitory role on methylation, while a negative score suggested an activating role. Finally, these raw scores were transformed into a comparable importance score for each gene using piecewise min-max scaling.

For interpreting the biological functions of the most impactful genes, we performed a pathway analysis using pre-computed KEGG enrichment results (KEGG 2021 Human^24^, via Enrichr^37^). Pathways with an adjusted p-value below 0.05 were considered significant. For visualization, we ranked genes by their absolute importance scores and annotated the top performers according to their membership in the most significantly enriched pathways.

### Statistics & Reproducibility

This is a computational study; all analyses were performed on pre-existing public datasets and no new biological samples were collected. No statistical method was used to predetermine sample size; and all variant-CpG pairs from the three meQTL cohorts that passed the pre-specified filters. No data were excluded from the analyses beyond these pre-specified criteria, namely the retention of variant-CpG pairs with p < 1e-5 and absolute methylation effect size > 0.5, and the requirement that sampled methylation blocks contain at least four CpG sites. The experiments were not randomized. The Investigators were not blinded to allocation during experiments and outcome assessment.

## Data availability

All the datasets and processing datasets code in this study have been deposited in Zenodo and are publicly available at https://doi.org/10.5281/zenodo.21386471 and https://doi.org/10.5281/zenodo.21386856. The raw data used in this study are available from public databases. The raw DNA methylation data used in this study are available in formats bigWig at GEO under accession code GSE186458 (https://www.ncbi.nlm.nih.gov/geo/query/acc.cgi?acc=GSE186458). The raw data for Ólafur et al. study^8^ are available at https://www.decode.com/summarydata/. The raw data for GTEx dataset^7^ are available at https://gtexportal.org/home/downloads/egtex/methylation. The raw data for EPIGEN resource are available at https://epicmeqtl.kcl.ac.uk/. The scRNA-seq data accession numbers used can be found in the Supplementary Table 3.

## Code availability

All the codes are publicly available at GitHub (https://github.com/FakeEnd/Melody) and available at Zenodo (https://doi.org/10.5281/zenodo.21386471). A separate GitHub repository contains code relevant to the analyses and results presented in the manuscript, located at https://github.com/FakeEnd/Melody_manuscript and available at Zenodo (https://doi.org/10.5281/zenodo.21386856). The developed web server is accessible via https://inner.wei-group.net/Melody/ and enables users to predict methylation levels across user-defined chromosomal regions (Supplementary Figure S9).

## Acknowledgments

We would like to thank the developers and maintainers of the public databases and software resources used in this study.

## Author Contributions

J.J., D.W. and J.Q. conceptualized the study, developed the methodology, implemented the model, performed the analyses, and wrote the original draft. W.G. and Y.L. plotted figures. S.C. built server. W.G., Y.L. and S.C. conducted the literature investigation and performed data analysis. Q.Z., S.W., R.S. and L.W. contributed to the writing, review, and editing of the manuscript and provided guidance and advice. R.S. and L.W. acquired funding and supervised the project. All authors discussed the results and contributed to the final manuscript.

## References

1. Moore, L. D., Le, T. & Fan, G. DNA methylation and its basic function. Neuropsychopharmacology 38, 23–38 (2013).

2. Smith, Z. D. & Meissner, A. DNA methylation: roles in mammalian development. Nat. Rev. Genet. 14, 204–220 (2013).

3. Koch, A. et al. Analysis of DNA methylation in cancer: location revisited. Nat. Rev. Clin. Oncol. 15, 459–466 (2018).

4. Robertson, K. D. DNA methylation and human disease. Nat. Rev. Genet. 6, 597–610 (2005).

5. Samblas, M., Milagro, F. I. & Martínez, A. DNA methylation markers in obesity, metabolic syndrome, and weight loss. Epigenetics 14, 421–444 (2019).

6. Bell, C. G. et al. DNA methylation aging clocks: challenges and recommendations. Genome Biol. 20, 249 (2019).

7. Oliva, M. et al. DNA methylation QTL mapping across diverse human tissues provides molecular links between genetic variation and complex traits. Nat. Genet. 55, 112–122 (2023).

8. Stefansson, O. A. et al. The correlation between CpG methylation and gene expression is driven by sequence variants. Nat. Genet. 56, 1624–1631 (2024).

9. Li, Z. et al. Applications of deep learning in understanding gene regulation. *Cell Rep*. Methods 3, 100384 (2023).

10. Sokolova, K., Chen, K. M., Hao, Y., Zhou, J. & Troyanskaya, O. G. Deep learning sequence models for transcriptional regulation. Annu. Rev. Genomics Hum. Genet. 25, 105–122 (2024).

11. Avsec, Ž., et al. AlphaGenome: advancing regulatory variant effect prediction with a unified DNA sequence model. bioRxiv (2025) doi:10.1101/2025.06.25.661532.

12. Avsec, Ž., et al. Effective gene expression prediction from sequence by integrating long-range interactions. Nat. Methods 18, 1196–1203 (2021).

13. Angermueller, C., Lee, H. J., Reik, W. & Stegle, O. DeepCpG: accurate prediction of single-cell DNA methylation states using deep learning. Genome Biol. 18, 67 (2017).

14. Zeng, H. & Gifford, D. K. Predicting the impact of non-coding variants on DNA methylation. Nucleic Acids Res. 45, e99 (2017).

15. Yu, Y. et al. iDNA-ABT: advanced deep learning model for detecting DNA methylation with adaptive features and transductive information maximization. Bioinformatics 37, 4603–4610 (2021).

16. Jin, J. et al. iDNA-ABF: multi-scale deep biological language learning model for the interpretable prediction of DNA methylations. Genome Biol. 23, 219 (2022).

17. Zhou, J., Weinberger, D. R. & Han, S. Deep learning predicts DNA methylation regulatory variants in specific brain cell types and enhances fine mapping for brain disorders. Sci. Adv. 11, eadn1870 (2025).

18. Loyfer, N. et al. A DNA methylation atlas of normal human cell types. Nature 613, 355–364 (2023).

19. Kathail, P. et al. Current genomic deep learning models display decreased performance in cell type-specific accessible regions. Genome Biol. 25, 202 (2024).

20. Villicaña, S. et al. Genetic impacts on DNA methylation help elucidate regulatory genomic processes. Genome Biol. 24, 176 (2023).

21. Novakovsky, G., Dexter, N., Libbrecht, M. W., Wasserman, W. W. & Mostafavi, S. Obtaining genetics insights from deep learning via explainable artificial intelligence. Nat. Rev. Genet. 24, 125–137 (2023).

22. Vierstra, J. et al. Global reference mapping of human transcription factor footprints. Nature 583, 729–736 (2020).

23. Monteagudo-Sánchez, A., Noordermeer, D. & Greenberg, M. V. C. The impact of DNA methylation on CTCF-mediated 3D genome organization. Nat. Struct. Mol. Biol. 31, 404–412 (2024).

24. Kanehisa, M., Furumichi, M., Sato, Y., Ishiguro-Watanabe, M. & Tanabe, M. KEGG: integrating viruses and cellular organisms. Nucleic Acids Res 49, D545– D551 (2021).

25. KEGG, Kyoto Encyclopedia of Genes and Genomes. (2001).

26. Schedel, J. et al. Targeting cathepsin L (CL) by specific ribozymes decreases CL protein synthesis and cartilage destruction in rheumatoid arthritis. Gene Ther 11, 1040–1047 (2004).

27. Iwata, Y., Mort, J. S., Tateishi, H. & Lee, E. R. Macrophage cathepsin L, a factor in the erosion of subchondral bone in rheumatoid arthritis. Arthritis Rheum 40, 499–509 (1997).

28. Hasegawa, T. et al. Identification of a novel arthritis-associated osteoclast precursor macrophage regulated by FoxM1. Nat Immunol 20, 1631–1643 (2019).

29. Mun, S. H. et al. Augmenting MNK1/2 activation by c-FMS proteolysis promotes osteoclastogenesis and arthritic bone erosion. Bone Res 9, 45 (2021).

30. Chen, K. M., Cofer, E. M., Zhou, J. & Troyanskaya, O. G. Selene: a PyTorch- based deep learning library for sequence data. Nat. Methods 16, 315–318 (2019).

31. Ronneberger, O., Fischer, P. & Brox, T. U-Net: Convolutional Networks for Biomedical Image Segmentation. *arXiv [cs.CV]* (2015) doi:10.48550/ARXIV.1505.04597.

32. Wolf, F. A., Angerer, P. & Theis, F. J. SCANPY: large-scale single-cell gene expression data analysis. Genome Biol 19, 15 (2018).

33. Hao, M. et al. Large-scale foundation model on single-cell transcriptomics. Nat Methods 21, 1481–1491 (2024).

34. Cui, H. et al. scGPT: toward building a foundation model for single-cell multi- omics using generative AI. Nat Methods 21, 1470–1480 (2024).

35. Zeng, Y. et al. CellFM: a large-scale foundation model pre-trained on transcriptomics of 100 million human cells. Nat Commun 16, 4679 (2025).

36. Perez, E., Strub, F., de Vries, H., Dumoulin, V. & Courville, A. FiLM: Visual reasoning with a general conditioning layer. *arXiv [cs.CV]* (2017) doi:10.48550/ARXIV.1709.07871.

37. Xie, Z. et al. Gene Set Knowledge Discovery with Enrichr. Curr Protoc 1, e90 (2021).

