## Supplementary information for "Melody: Decoding the Sequence Determinants of Locus Specific DNA Methylation Across Human Tissues"

[#]These authors contributed equally

*** Correspondence:**

### 1. Supplementary Tables

**Supplementary Table 1** lists the 39 cell-type–specific bigWig tracks used as references. The left column gives the abbreviated cell-type names (also used in Supplementary Table 2), and the right column shows the corresponding bigWig file names.

| Cell type (abbreviation) | bigWig file name |
|---|---|
| Adipocytes | GSM5652176_Adipocytes-Z000000T7.hg38.bigwig |
| Aorta_Endothel | GSM5652179_Aorta-Endothel-Z00000422.hg38.bigwig |
| Kidney_Tubular_Endothel | GSM5652189_Kidney-Tubular-Endothel-Z0000042R.hg38.bigwig |
| Colon_Fibroblasts | GSM5652198_Colon-Fibroblasts-Z0000042A.hg38.bigwig |
| Heart_Fibroblasts | GSM5652200_Heart-Fibroblasts-Z0000043R.hg38.bigwig |
| Dermal_Fibroblasts | GSM5652204_Dermal-Fibroblasts-Z00000423.hg38.bigwig |
| Skeletal_Muscle | GSM5652205_Skeletal-Muscle-Z00000427.hg38.bigwig |
| Aorta_SmoothMuscle | GSM5652207_Aorta-Smooth-Muscle-Z0000041U.hg38.bigwig |
| Heart_Cardiomyocyte | GSM5652215_Heart-Cardiomyocyte-Z0000044P.hg38.bigwig |
| Bone_Osteoblasts | GSM5652218_Bone-Osteoblasts-Z0000042Z.hg38.bigwig |
| Oligodendrocytes | GSM5652219_Oligodendrocytes-Z000000TK.hg38.bigwig |
| Cortex_Neuron | GSM5652223_Cortex-Neuron-Z000000TF.hg38.bigwig |
| Liver_Hepatocytes | GSM5652233_Liver-Hepatocytes-Z000000R3.hg38.bigwig |
| Pancreas_Duct | GSM5652239_Pancreas-Duct-Z0000043T.hg38.bigwig |
| Pancreas_Acinar | GSM5652243_Pancreas-Acinar-Z000000QX.hg38.bigwig |
| Pancreas_Delta | GSM5652247_Pancreas-Delta-Z00000451.hg38.bigwig |
| Pancreas_Beta | GSM5652250_Pancreas-Beta-Z00000452.hg38.bigwig |
| Pancreas_Alpha | GSM5652253_Pancreas-Alpha-Z00000453.hg38.bigwig |
| Thyroid_Epithelial | GSM5652264_Thyroid-Epithelial-Z0000042S.hg38.bigwig |
| Fallopian_Epithelial | GSM5652267_Fallopian-Epithelial-Z000000Q7.hg38.bigwig |
| Ovary_Epithelial | GSM5652270_Ovary-Epithelial-Z000000QT.hg38.bigwig |

| | |
|---|---|
| BM_Erythro_Prog | GSM5652274_Bone_marrow-Erythrocyte_progenitors-Z000000RF.hg38.bigwig |
| Blood_T_CD3 | GSM5652277_Blood-T-CD3-Z000000TV.hg38.bigwig |
| Blood_NK | GSM5652299_Blood-NK-Z000000TM.hg38.bigwig |
| Blood_Monocytes | GSM5652302_Blood-Monocytes-Z000000TP.hg38.bigwig |
| Blood_Granulocytes | GSM5652313_Blood-Granulocytes-Z000000TZ.hg38.bigwig |
| Blood_B | GSM5652317_Blood-B-Z000000UB.hg38.bigwig |
| Epidermal_Keratinocytes | GSM5652321_Epidermal-Keratinocytes-Z00000424.hg38.bigwig |
| Tonsil_Palatine_Epi | GSM5652322_Tonsil-Palatine-Epithelial-Z000000QF.hg38.bigwig |
| Lung_Bronchus_Epi | GSM5652335_Lung-Bronchus-Epithelial-Z000000QD.hg38.bigwig |
| Prostate_Epithelial | GSM5652338_Prostate-Epithelial-Z000000RV.hg38.bigwig |
| Bladder_Epithelial | GSM5652342_Bladder-Epithelial-Z000000QM.hg38.bigwig |
| Breast_Luminal_Epi | GSM5652347_Breast-Luminal-Epithelial-Z000000V2.hg38.bigwig |
| Breast_Basal_Epi | GSM5652350_Breast-Basal-Epithelial-Z000000V6.hg38.bigwig |
| Lung_Alveolar_Epi | GSM5652354_Lung-Alveolar-Epithelial-Z000000T1.hg38.bigwig |
| Gallbladder_Epi | GSM5652358_Gallbladder-Epithelial-Z00000432.hg38.bigwig |
| GastricFundus_Epi | GSM5652359_Gastric-fundus-Epithelial-Z000000RX.hg38.bigwig |
| Colon_Right_Epi | GSM5652370_Colon-Right-Epithelial-Z000000V0.hg38.bigwig |
| SmallInt_Epi | GSM5652378_Small-int-Epithelial-Z0000042V.hg38.bigwig |

**Supplementary Table 2.** The meQTL datasets and the corresponding reference cell-type tracks used in the evaluation.

| meQTLs type | meQTLs dataset | Corresponding cell-type tracks |
|---|---|---|
| Olafur | CPG_units | Bone marrow erythrocyte progenitors; blood T cells (CD3$^{+}$); blood NK cells; blood monocytes; blood granulocytes; blood B cells |
| | MDSs | Bone marrow erythrocyte progenitors; blood T cells (CD3$^{+}$); blood NK cells; blood monocytes; blood granulocytes; blood B cells |
| GTEx Datasets | GTEx_WholeBlood | Bone marrow erythrocyte progenitors; blood T cells (CD3$^{+}$); blood NK cells; blood monocytes; blood granulocytes; blood B cells |
| | GTEx_BreastMammaryTissue | Breast luminal epithelial; breast basal epithelial; adipocytes |
| | GTEx_ColonTransverse | Right colon epithelial; colon fibroblasts; small-intestinal epithelial |
| | GTEx_KidneyCortex | Kidney tubular endothelial |
| | GTEx_Lung | Bronchial epithelial; alveolar epithelial |
| | GTEx_MuscleSkeletal | Skeletal muscle; aorta smooth muscle |
| | GTEx_Ovary | Ovary epithelial; fallopian epithelial |
| | GTEx_Prostate | Prostate epithelial |
| EPIGEN | skin | Epidermal keratinocytes; dermal fibroblasts |

| | |
|---|---|
| EPIC | Bone marrow erythrocyte progenitors; blood T cells (CD3$^+$); blood NK cells; blood monocytes; blood granulocytes; blood B cells |

**Supplementary Table 3.** The accession numbers corresponding to the scRNA-seq data for the cell types used.

| **Cell types** | **Project ID or Pubmed ID** |
|---|---|
| Adipocytes | GSE163830 |
| Aorta-Endothel | GSE155468 |
| Aorta-Smooth-Muscle | 33017217 |
| Bladder-Epithelial | GSE146137 |
| Blood-B | GSE182270 |
| Blood-Granulocytes | GSE124494 |
| Blood-Monocytes | GSE142392, GSE167029 |
| Blood-NK | GSE158349 |
| Blood-T-CD3 | GSE138720 |
| Bone-Osteoblasts | GSE143753 |
| Bone_marrow-Erythrocyte_progenitors | GSE190067 |
| Breast-Basal-Epithelial | GSE164898 |
| Breast-Luminal-Epithelial | GSE145326 |
| Colon-Fibroblasts | GSE125527 |
| Colon-Right-Epithelial | GSE188711 |
| Cortex-Neuron | GSE175719 |
| Dermal-Fibroblasts | GSE132802 |
| Epidermal-Keratinocytes | GSE150672 |
| Fallopian-Epithelial | GSE118127 |

| | |
|---|---|
| Gallbladder-Epithelial | GSE175502 |
| Gastric-fundus-Epithelial | GSE58557 |
| Heart-Cardiomyocyte | GSE148586, GSE161153 |
| Heart-Fibroblasts | GSE145154 |
| Kidney-Tubular-Endothel | GSE131685 |
| Liver-Hepatocytes | GSE136103 |
| Lung-Alveolar-Epithelial | GSE168191 |
| Lung-Bronchus-Epithelial | GSE121600 |
| Oligodendrocytes | GSE118257 |
| Ovary-Epithelial | GSE118127 |
| Pancreas-Acinar | GSE153834 |
| Pancreas-Alpha | GSE159556 |
| Pancreas-Beta | GSE114297, GSE139535, GSE143783 |
| Pancreas-Delta | GSE153855 |
| Pancreas-Duct | GSE153855 |
| Prostate-Epithelial | GSE130318, GSE142489 |
| Skeletal-Muscle | GSE130646 |
| Small-int-Epithelial | GSE185224 |
| Thyroid-Epithelial | GSE184362 |
| Tonsil-Palatine-Epithelial | DISCO-E-MTAB-9005 |

#### 2. Supplementary Figures

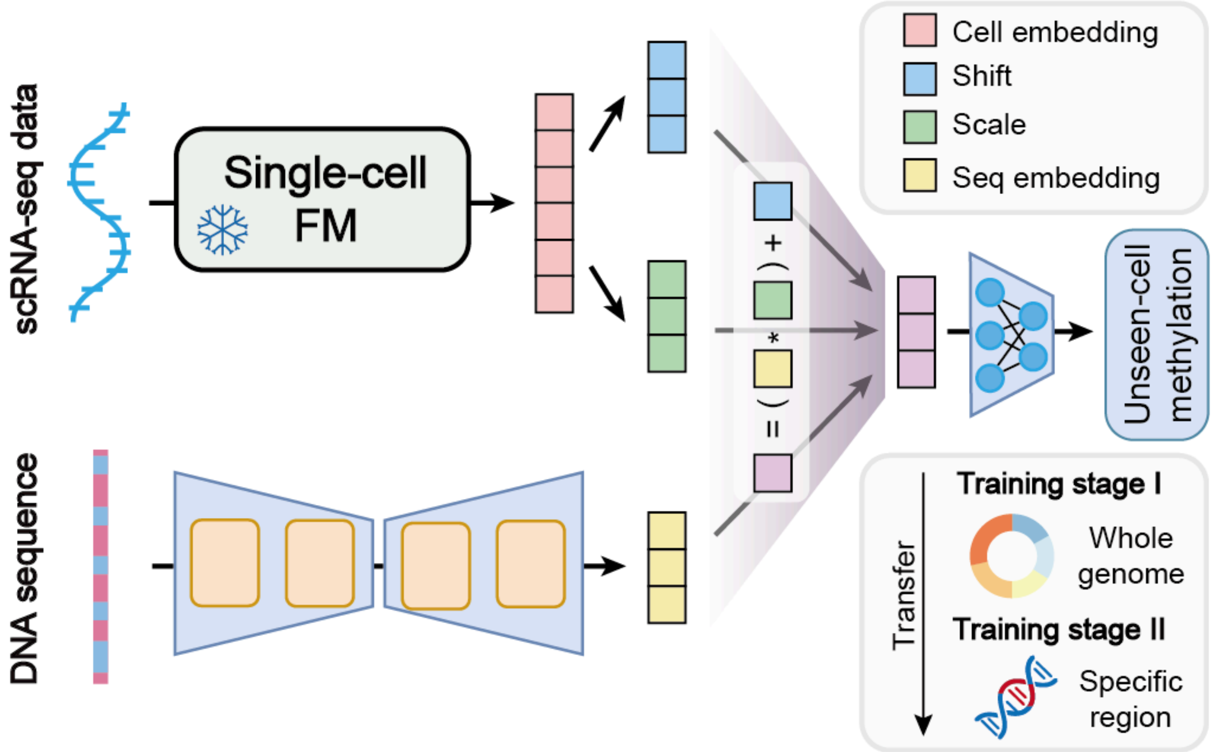


**Supplementary Figure S1**. Melody-G is designed for methylation level prediction in unseen cell types. It utilizes DNA sequences alongside scRNA-seq data encoded by a single-cell foundation model (FM). By fusing these multimodal inputs via feature-wise linear modulation and employing a two-stage training strategy, the model achieves cell-type-specific methylation profiling for previously unobserved cell types.

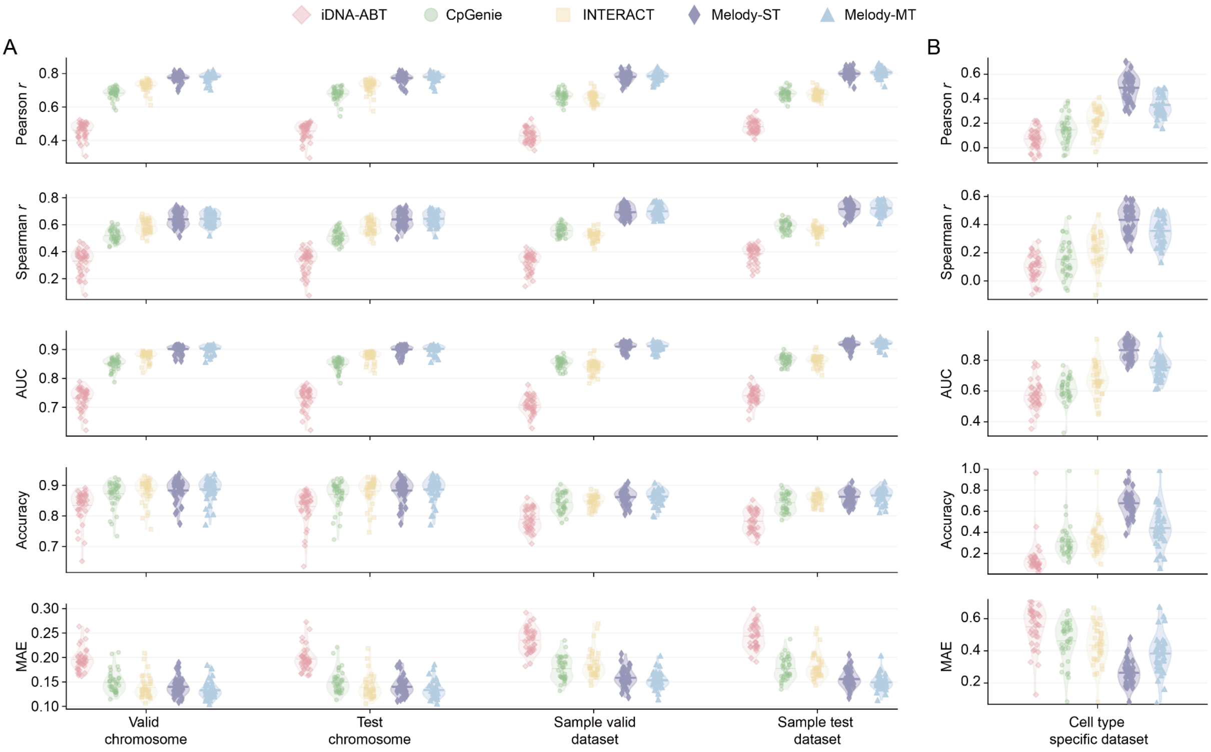


**Supplementary Figure S2**. Comprehensive benchmarking of Melody-MT and Melody-ST against the three leading methylation predictors (iDNA-ABT, CpGenie, INTERACT) across all evaluation splits and metrics. (A) Performance on four genome-wide splits (validation chromosome, test chromosome, hypomethylated sampling validation set, hypomethylated sampling test set) using five complementary metrics: Pearson correlation, Spearman rank correlation, AUC, Accuracy, and MAE. Each dot represents one of 39 cell-type datasets. (B) Performance on cell-type-specific regions (n = 38 cell-type datasets). Across every split × metric combination, both Melody variants outperform all three baselines, with the gap widening substantially on the hypomethylated sampling sets and on cell-type-specific regions, where Melody-ST achieves the strongest performance.

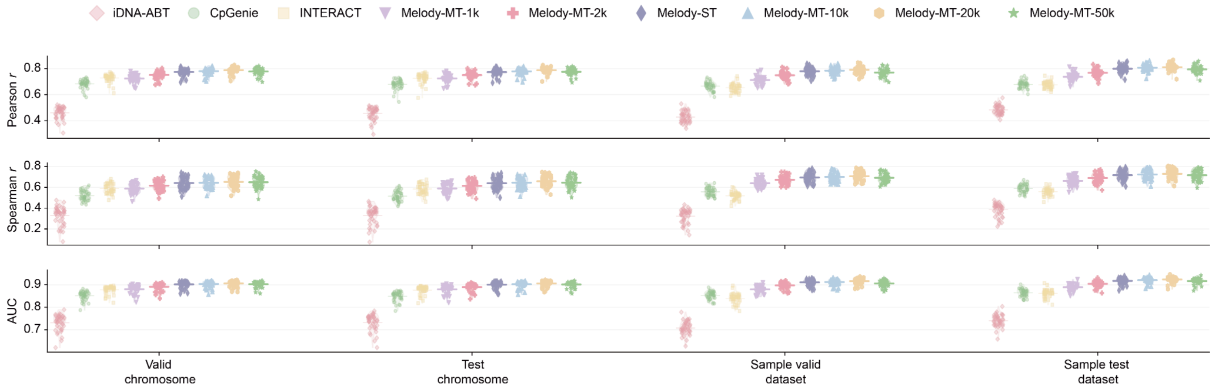


**Supplementary Figure S3**. Contribution of sequence window length to Melody's advantage over short-window baselines. Matched-architecture Melody-MT variants at 1-kb, 2-kb, 10-kb, 20-kb, and 50-kb input windows, together with Melody-ST, are compared against iDNA-ABT (41 bp), CpGenie (1 kb), and INTERACT (2 kb) on three continuous metrics (Pearson, Spearman, AUC) across four genome-wide splits. Each point is one of 39 cell-type datasets. At matched short input windows, Melody-MT-1k performs comparably to INTERACT (2 kb); extending the window to 10 kb produces the main performance lift, with 20-kb and 50-kb variants offering no further gain on genome-wide metrics. This decomposition indicates that both the U-Net architecture and the 10-kb receptive field contribute to Melody's final performance, with long-range context being the dominant source of gain.

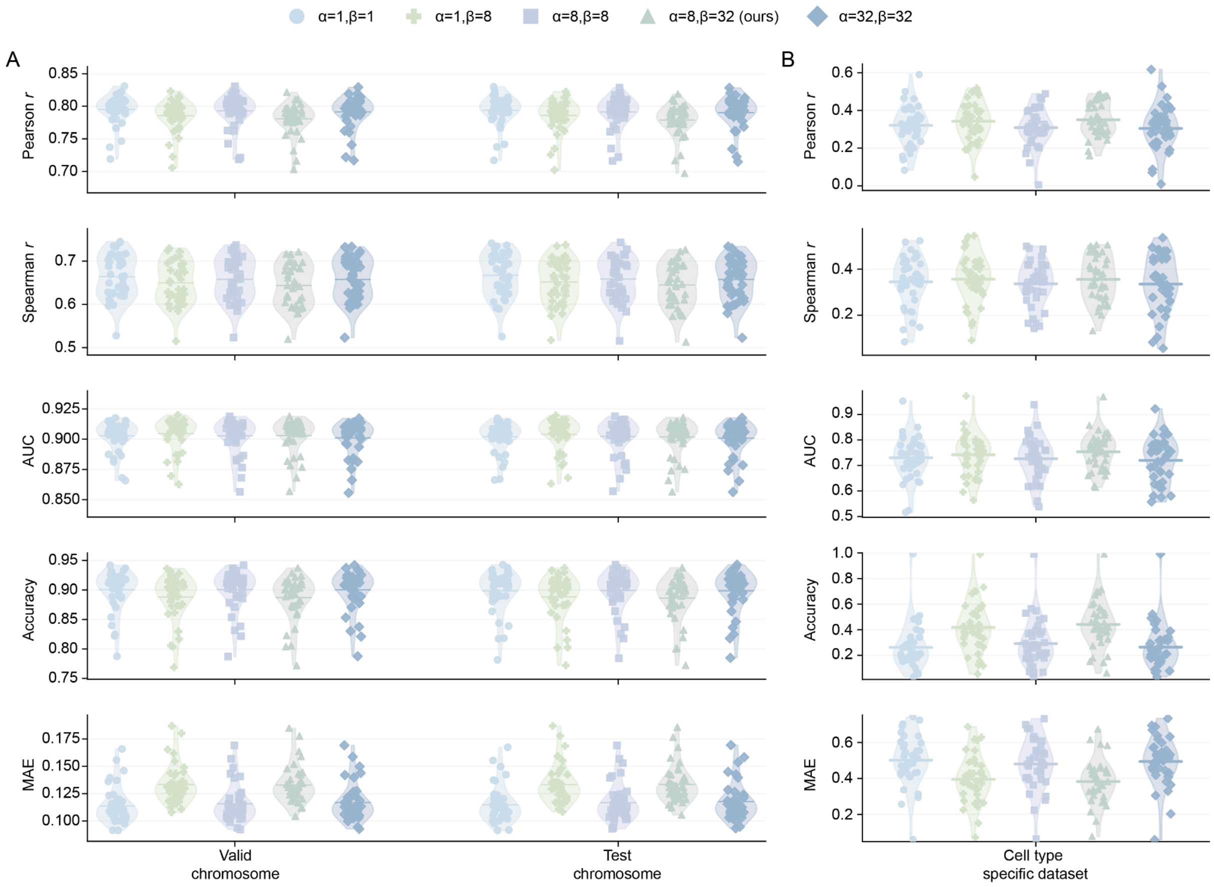


**Supplementary Figure S4**. Sensitivity of Melody-MT performance to the CpG / low-methylation-CpG loss weights (α, β). Five configurations were trained to 1,000,000 steps with identical settings: α=1,β=1 (unweighted), α=1,β=8, α=8,β=8, α=8,β=32 (Melody default), and α=32,β=32. (A) Genome-wide performance on the validation and test chromosomes across five metrics (n = 39 cell-type datasets); the five configurations show nearly identical distributions (Pearson spread within ±2%, AUC within ±0.4%), confirming that genome-wide prediction is robust to the choice of (α, β). (B) Performance on cell-type-specific regions (n = 38 cell-type datasets). The default α=8,β=32 configuration achieves the highest Pearson/Spearman/AUC/Accuracy and the lowest MAE, with the unweighted baseline (α=1,β=1) showing approximately 24% higher MAE on cell type specific dataset, supporting our choice of weighting scheme for cell-type-specific hypomethylated regulatory regions.

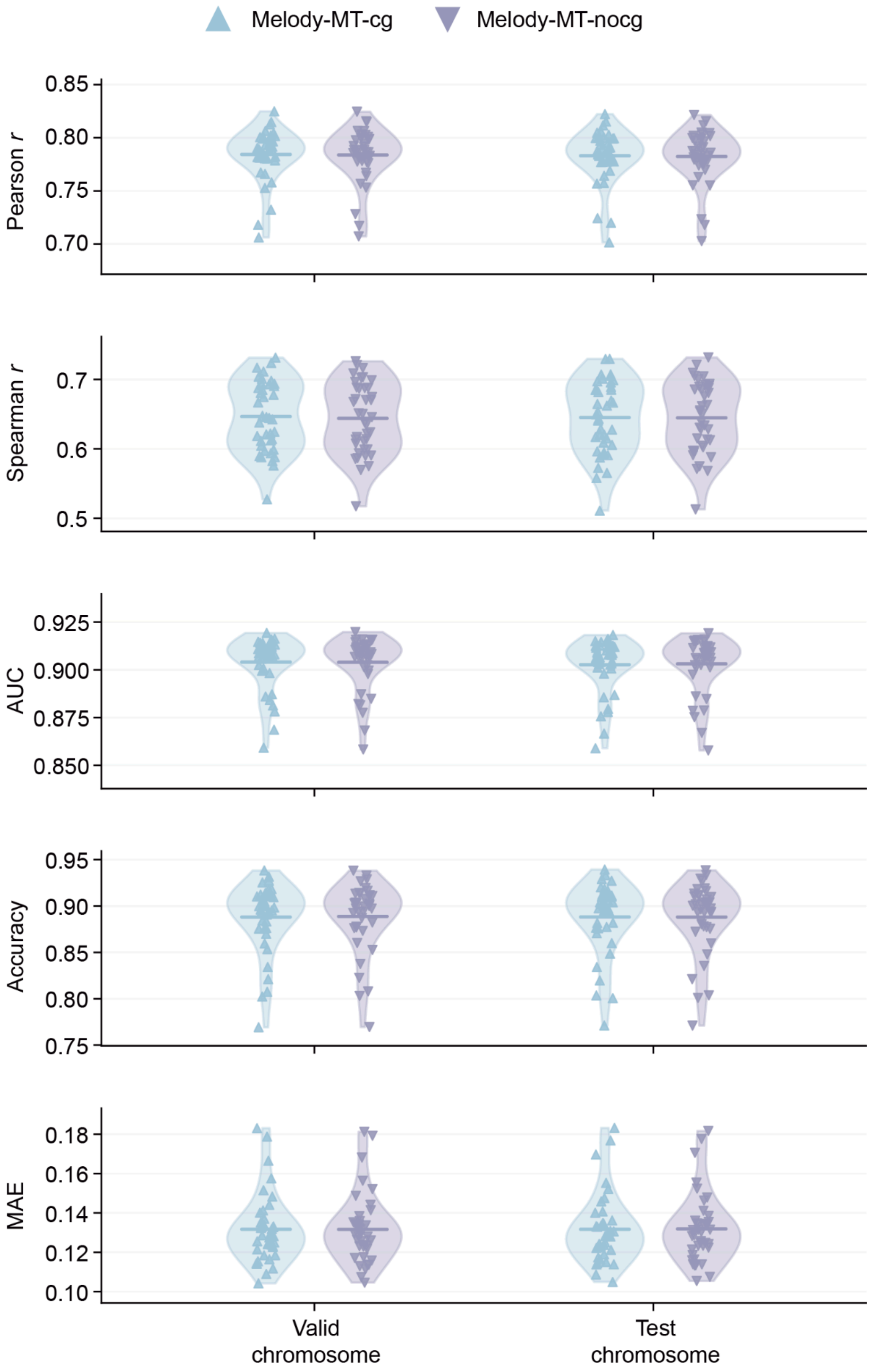


**Supplementary Figure S5.** Effect of the CpG-count auxiliary loss on Melody-MT performance. Violin plots compare Melody-MT trained with (cg) versus without (nocg) the CpG-count auxiliary head across five continuous evaluation metrics (Pearson, Spearman, AUC, Accuracy, MAE) on the validation and test chromosomes. Each point is one of the 39 cell types. The two variants yield essentially overlapping distributions (all differences within ±1%), demonstrating that removal of the CpG-count loss does not degrade methylation prediction and that the primary model performance is not driven by CpG-counting shortcut behavior.

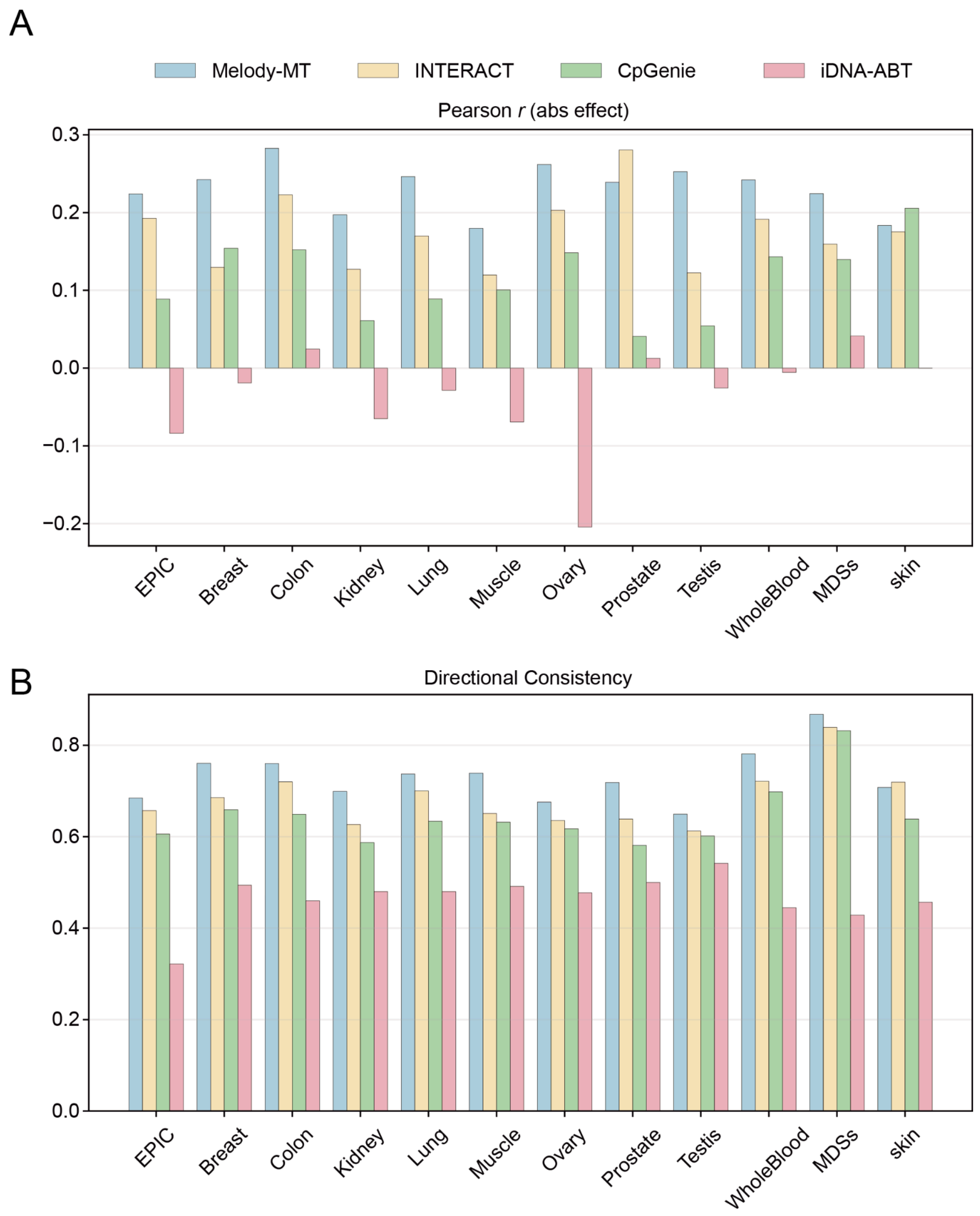


**Supplementary Figure S6**. Direction- and magnitude-based evaluation of meQTL effect prediction. (A) Pearson correlation between predicted and observed absolute meQTL effect sizes per dataset, which isolates magnitude agreement from directional consistency. Melody-MT achieves positive correlations across all 13 datasets, whereas iDNA-ABT frequently yields negative or near-zero correlations, indicating that Melody's magnitude prediction is directional-robust. (B) Directional consistency per dataset (fraction of SNP–CpG pairs whose predicted effect sign matches the observed sign). Melody-MT achieves the highest directional consistency on every dataset, reaching 88% on MDSs and 78% on GTEx WholeBlood, consistently above the 50% random baseline. Together these panels demonstrate that Melody's meQTL prediction performance reflects both accurate directional and magnitude estimation rather than direction alone.

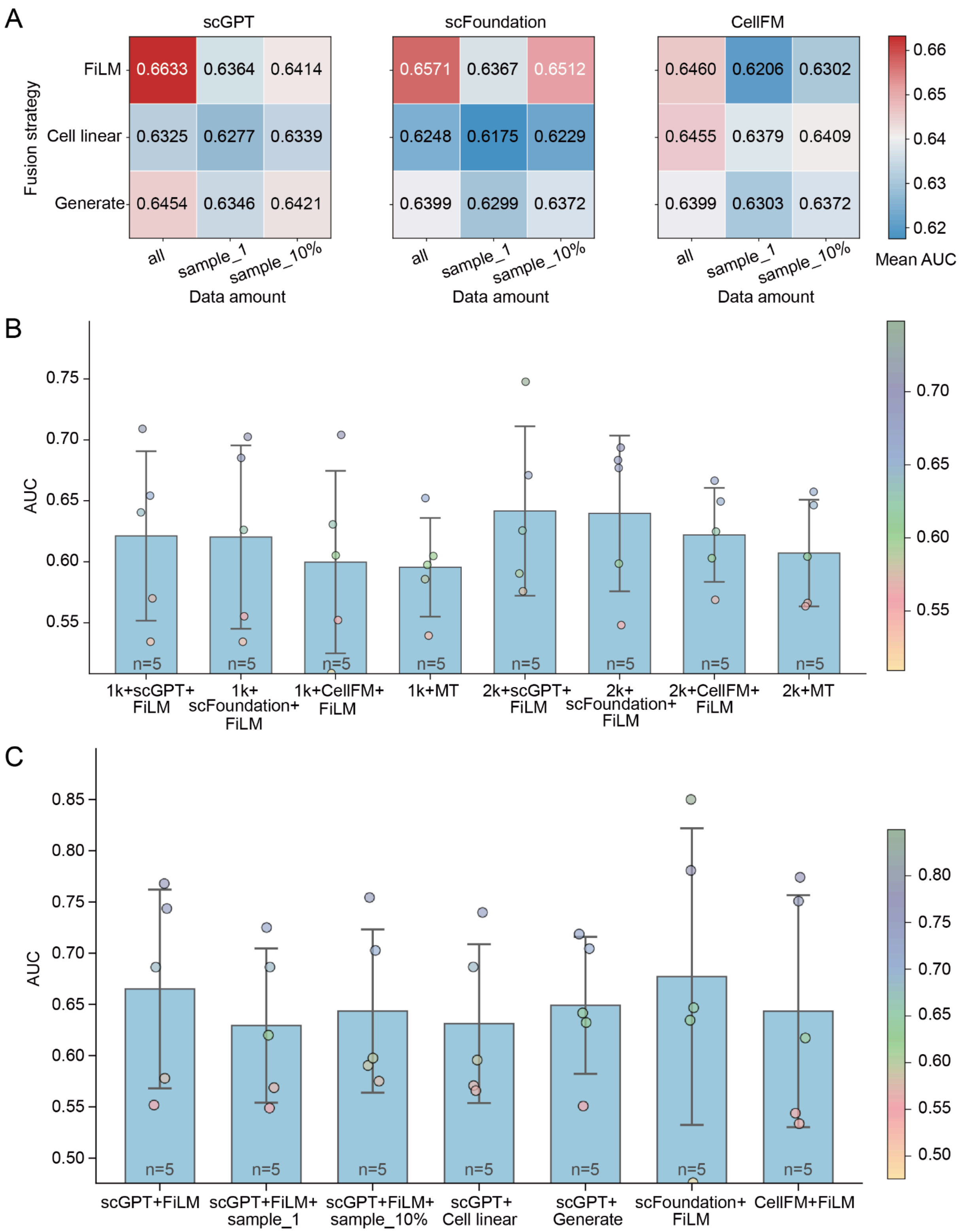


**Supplementary Figure S7**. Systematic ablation of Melody-G components. (A) Full 3×3×3 heatmap (foundation models × fusion strategies × sampling settings; mean AUC across five unseen cell types). FiLM shows the best overall performance, with the strongest and most consistent gains for scFoundation and scGPT, while the relative advantage is weaker for CellFM under reduced sampling settings. (B) Performance of Melody-G under matched 1-kb and 2-kb input windows using three foundation-model backbones (scGPT, scFoundation, CellFM) with FiLM fusion, compared against the corresponding Melody-MT baseline at the same window size. At both window scales, Melody-G variants outperform the MT baseline (e.g., at 1 kb, scGPT+FiLM 0.621 vs MT 0.596), confirming that the transcriptomic embeddings contribute beyond sequence length alone. (C) Performance under the newly defined unseen cell type split across foundation models and fusion strategies. For (B) and (C), scatter points represent the performance distribution across n=5 unseen cell lines. Bars show the mean AUC across five unseen cell types; error bars indicate the standard deviation; points represent individual unseen cell types.

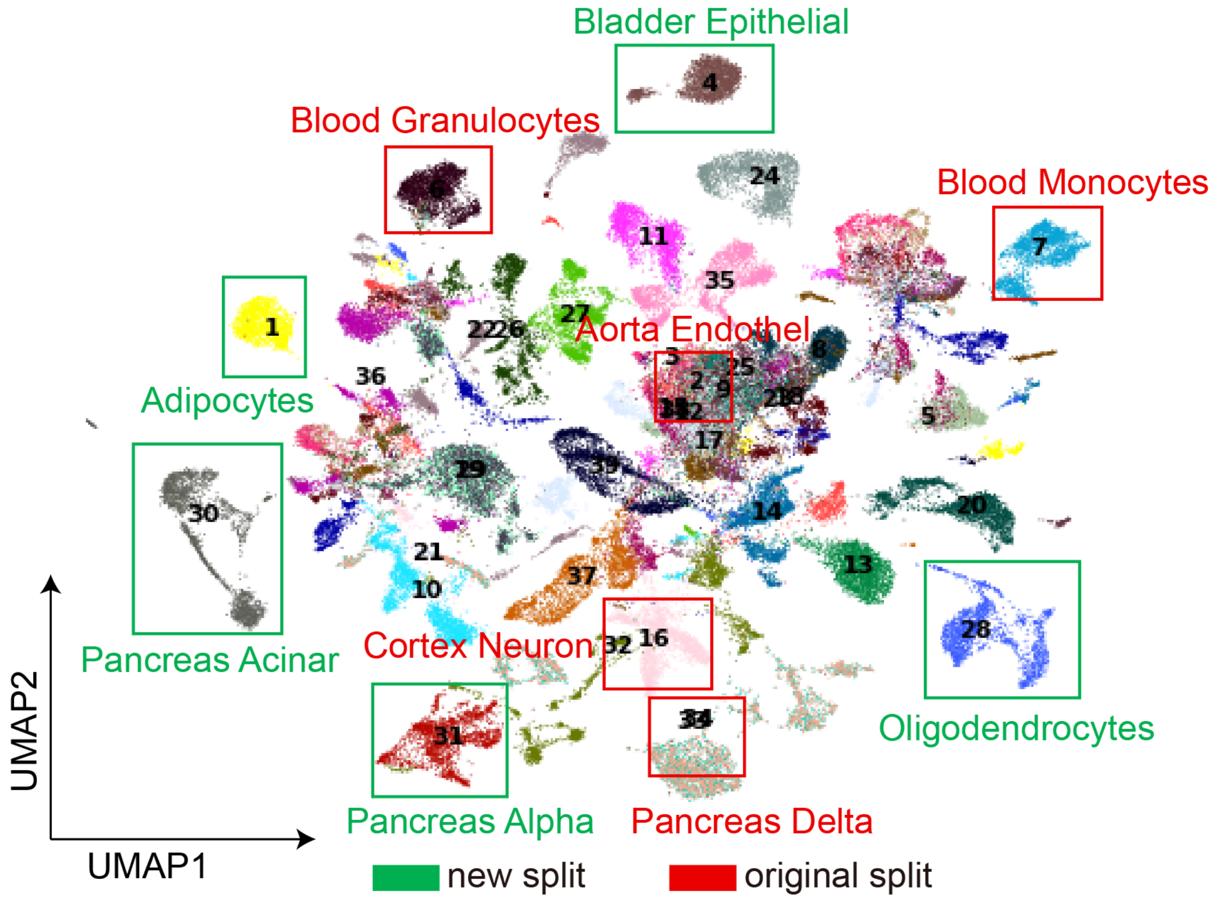


**Supplementary Figure S8**. UMAP of the 39 cell types based on the original scRNA-seq expression matrix. The UMAP was generated after normalization, log transformation, highly variable gene selection, PCA, and neighborhood graph construction. Red boxes indicate the five cell types used in the original held out split, namely Aorta Endothel, Blood Granulocytes, Blood Monocytes, Cortex Neuron, and Pancreas Delta. Green boxes indicate the five cell types used in the newly defined held out split, namely Pancreas Acinar, Pancreas Alpha, Oligodendrocytes, Adipocytes, and Bladder Epithelial. Compared with the original split, the newly selected held out cell types appear more clearly separated from the main transcriptomic cluster and were therefore used as a more stringent test of cross cell type generalization.

**Label to cell type mapping for Supplementary Figure S8**: 1, Adipocytes; 2, Aorta Endothel; 3, Aorta Smooth Muscle; 4, Bladder Epithelial; 5, Blood B; 6, Blood Granulocytes; 7, Blood Monocytes; 8, Blood NK; 9, Blood T CD3; 10, Bone Osteoblasts; 11, Bone marrow Erythrocyte progenitors; 12, Breast Basal Epithelial; 13, Breast Luminal Epithelial; 14, Colon Fibroblasts; 15, Colon Right Epithelial; 16, Cortex Neuron; 17, Dermal Fibroblasts; 18, Epidermal Keratinocytes; 19, Fallopian Epithelial; 20, Gallbladder Epithelial; 21, Gastric fundus Epithelial; 22, Heart Cardiomyocyte; 23, Heart Fibroblasts; 24, Kidney Tubular Endothel; 25, Liver Hepatocytes; 26, Lung Alveolar Epithelial; 27, Lung Bronchus Epithelial; 28, Oligodendrocytes; 29, Ovary Epithelial; 30, Pancreas Acinar; 31, Pancreas Alpha; 32, Pancreas Beta; 33,

Pancreas Delta; 34, Pancreas Duct; 35, Prostate Epithelial; 36, Skeletal Muscle; 37, Small intestine Epithelial; 38, Thyroid Epithelial; 39, Tonsil Palatine Epithelial.

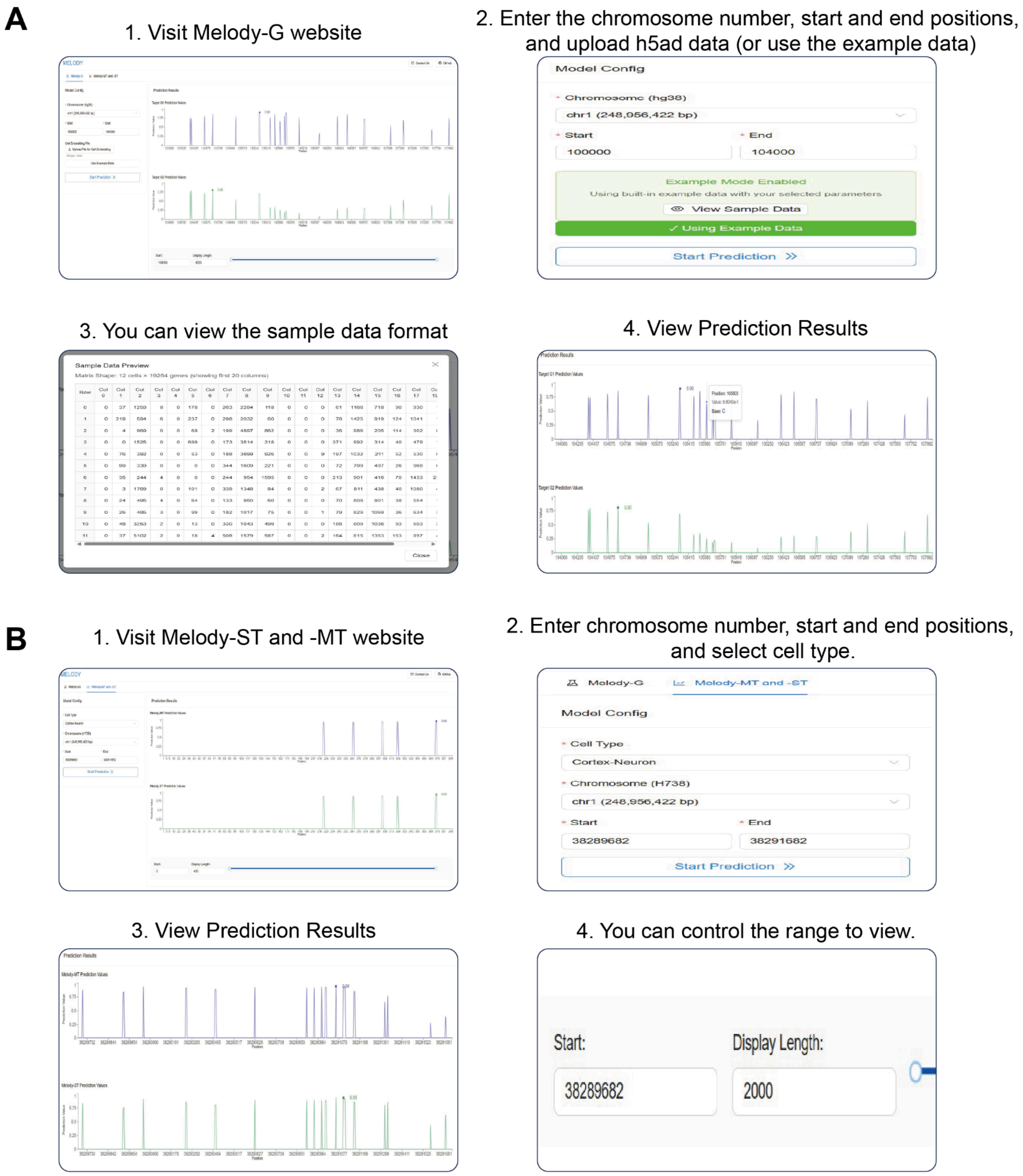


**Supplementary Figure S9. Melody web interface workflow example operation guide.** (A) Melody-G workflow. Users define the prediction scope by selecting chromosomes and specifying start and end coordinates. The platform accommodates user-uploaded scRNA-seq data in .h5ad format containing a cell by gene matrix or allows the utilization of preloaded example datasets. The panel displays results generated by both Melody-G1 and Melody-G2. (B) Workflow for Melody-ST and Melody-MT. Users specify target cell types and chromosomal intervals for analysis. The panel presents predictive outcomes derived from the Melody-ST and Melody-MT models.
